# Loss of *Grem1*-articular cartilage progenitor cells causes osteoarthritis

**DOI:** 10.1101/2023.03.29.534651

**Authors:** Jia Q. Ng, Toghrul H. Jafarov, Christopher B. Little, Tongtong Wang, Abdullah Ali, Yan Ma, Georgette A Radford, Laura Vrbanac, Mari Ichinose, Samuel Whittle, David Hunter, Tamsin RM Lannagan, Nobumi Suzuki, Jarrad M. Goyne, Hiroki Kobayashi, Timothy C. Wang, David Haynes, Danijela Menicanin, Stan Gronthos, Daniel L. Worthley, Susan L. Woods, Siddhartha Mukherjee

## Abstract

Osteoarthritis (OA), which carries an enormous disease burden across the world, is characterised by irreversible degeneration of articular cartilage (AC), and subsequently bone. The cellular cause of OA is unknown. Here, using lineage tracing in mice, we show that the BMP-antagonist *Gremlin 1* (*Grem1*) marks a novel chondrogenic progenitor (CP) cell population in the articular surface that generates joint cartilage and subchondral bone during development and adulthood. Notably, this CP population is depleted in injury-induced OA, and with age. OA is also induced by toxin-mediated ablation of *Grem1* CP cells in young mice. Transcriptomic analysis and functional modelling in mice revealed articular surface *Grem1*-lineage cells are dependent on *Foxo1*; ablation of *Foxo1* in *Grem1*-lineage cells led to early OA. This analysis identified FGFR3 signalling as a therapeutic target, and injection of its activator, FGF18, caused proliferation of *Grem1*-lineage CP cells, increased cartilage thickness, and reduced OA pathology. We propose that OA arises from the loss of CP cells at the articular surface secondary to an imbalance in progenitor cell homeostasis and present a new progenitor population as a locus for OA therapy.

## INTRODUCTION

OA is the most common articular disease of the developed world, affecting 10 percent of people over the age of 60 (Pereira et al., 2011). All tissues of the joint are affected including loss of AC, subchondral bone remodelling, osteophyte formation, synovial inflammation and joint capsule fibrosis, accompanied by instability, pain and disability (Madisen et al., 2010), necessitating joint replacement in 8-11% of patients with knee OA (Hunter and Bierma-Zeinstra, 2019). There is no cure for OA and existing treatments involve pain management and lifestyle modification (Maiese, 2016). There are several mouse models that induce OA via the surgical destabilization of medial meniscus (DMM) or collagenase-induced injury to destabilize tendons and ligaments (collagenase-induced OA, or CIOA). However, these models are descriptive, and the cellular pathophysiology of OA remains unknown.

Articular cartilage is formed by chondrocytes that secrete a rich extracellular matrix (ECM) with high proteoglycan content, to allow effective movement between two bones (Gannon et al., 2015; Helminen et al., 2000). Unlike the growth plate (GP) cartilage, AC is a permanent tissue that requires the support of self-renewing progenitor cells to repopulate resident chondrocytes (Creamer and Hochberg, 1997; Felson et al., 2000; Gannon et al., 2015; Harrison et al., 1953; Hashimoto et al., 2008; Li et al., 2017). AC has limited regenerative capacity upon injury, mechanical stress or in old age, with AC loss being a key feature of OA (Gannon et al., 2015; Kraus et al., 2015; Murphy et al., 2020).

Several groups, including ours, have described novel skeletal stem cells (Bianco and Robey, 2015) (SSC) based on immunophenotype (Chan et al., 2015) or lineage tracing (Worthley et al., 2015) that give rise to bone, cartilage and stroma but not fat lineages. Independently, a bone-fat (but not cartilage) progenitor cell population, marked by the expression of the *Leptin Receptor* (*Lepr*), was identified within the marrow (Zhou et al., 2014). The bone-cartilage-stromal progenitors in the growth plate and the bone-fat progenitors in the marrow express different markers (*Grem1* and *Lepr,* respectively) and have different fates during development and repair. Our initial study (Worthley et al., 2015) focused on *Grem1*-lineage tissue-resident SSCs in the GP of mice. A subsequent study found that tissue-resident SSC can be activated to make AC using microfractures in conjunction with BMP and VEGF signalling; however, the location of these AC-forming SSCs remained unknown, and these stimuli appeared to generate only cartilage, not subchondral bone (Murphy et al., 2020). Here, we show that a unique population of *Grem1*-lineage chondrogenic progenitor (CP) cells, distinct from GP-resident SCCs, resides on the articular surface, and generates AC (and, in later stages, subchondral bone). We focused on the fate and function of these articular surface CP cells during aging, and upon OA-inducing injury. We find that during aging, and in two independent models of OA-inducing injury (DMM and CIOA), this CP *Grem1*-lineage is lost through apoptosis. Toxin-mediated ablation of these CPs in young mice also caused OA. Single cell RNA sequencing (scRNAseq) of *Grem1* expressing cells at the articular surface revealed distinct molecular features of *Grem1*-lineage cells and FGFR3 agonists as potential therapeutic targets for *Grem1*-lineage CP cell maintenance and expansion. The FGFR3 ligand, FGF18 (the active agent in Sprifermin) is currently in human clinical trial for OA treatment (Hochberg et al., 2019). Injection of FGF18 into injured joints increased the number of *Grem1*-lineage CP cells (but not mitosis in hypertrophic chondrocytes) and ameliorated OA pathology. Our study thus identifies a novel, previously unknown cellular locus for OA pathophysiology and therapy, and posits that OA is a disease caused by CP cell loss at the articular surface.

## RESULTS

### *Grem1* CP cells are depleted in OA

Focusing our studies on the knee joint of adult mice, we observed two anatomically distinct populations in the AC and GP marked by *Grem1-*lineage tracing (**Figure 1A**). Physically these were separated in the femur by 800-1900 microns. As such, we first determined if the articular *Grem1*-lineage cells were affected under OA disease conditions. OA is predisposed by injuries that destabilise the periarticular tissues of a joint (Haq et al., 2003). Here we used two models of induced OA in mice, involving surgical DMM or collagenase VII degradation of intra-articular stabilising ligaments (CIOA), to examine the fate of *Grem1* lineage cells, *Lepr* bone marrow derived-mesenchymal stem cells (MSC) and articular chondrocytes marked by *Acan* in OA. *Rosa-TdTomato* reporter mice crossed with *Grem1-creERT*, *Acan-creERT* and *Lepr-cre* (a constitutive *Cre* line), henceforth termed *Grem1-TdT*, *Acan-TdT* and *Lepr-TdT* mice respectively, labelled *Grem1-*, *Acan-* and *Lepr-* lineages. The inducible Cre lines, *Acan-TdT* and *Grem1-TdT,* were administered tamoxifen at 8 to 10 weeks of age, with DMM surgery (**Figure S1A**) performed 2 weeks later. *Lepr-TdT* mice of similar age had DMM surgery for comparison (**Figure 1B**). DMM surgery results in a significant decrease in proliferating cells in both the superficial and non-calcified zones of the AC (**Figure S1B**). OA pathology was confirmed by loss of proteoglycans, surface fibrillation (**Figure 1C**) and osteophyte formation (**Figure 1D**). Quantification of the total percentage of *Acan*, *Grem1* and *Lepr* traced AC cells at the site of proteoglycan loss showed a significant decrease in the *Grem1*-lineage population only (**Figure 1E and 1F**). This suggested Grem1 AC chondrocytes may be important in preventing the progression of OA by secretion of proteoglycans, a process critical to maintaining AC integrity and protection from daily mechanical insult. The absence of lineage-traced cells in the AC of *Lepr-TdT* mice is consistent with a prior study (Zhou et al., 2014) and suggested that the primary role of the *Lepr*-lineage is in the haematopoietic niche in diaphyseal bone marrow (Ding et al., 2012; Mendez-Ferrer et al., 2010). Notably, the persistence of *Acan* AC chondrocytes in DMM animals suggests that the initial stage of OA is not due to total chondrocyte loss, but rather is specific to the loss of the *Grem1*-lineage CP population. In the CIOA model (**Figure 1G**), we observed more severe OA pathology as expected (Botter et al., 2008). As with DMM, *Grem1-TdT* CIOA mice exhibited significant loss of *Grem1*-lineage CP cells through apoptosis, decreased AC thickness and increased OA pathology (Glasson et al., 2010) compared to PBS injected animals (**Figure 1H-K and S1C**). This suggested *Grem1*-lineage articular surface CP cells may be important in the pathophysiology of OA.

**Figure 1.**
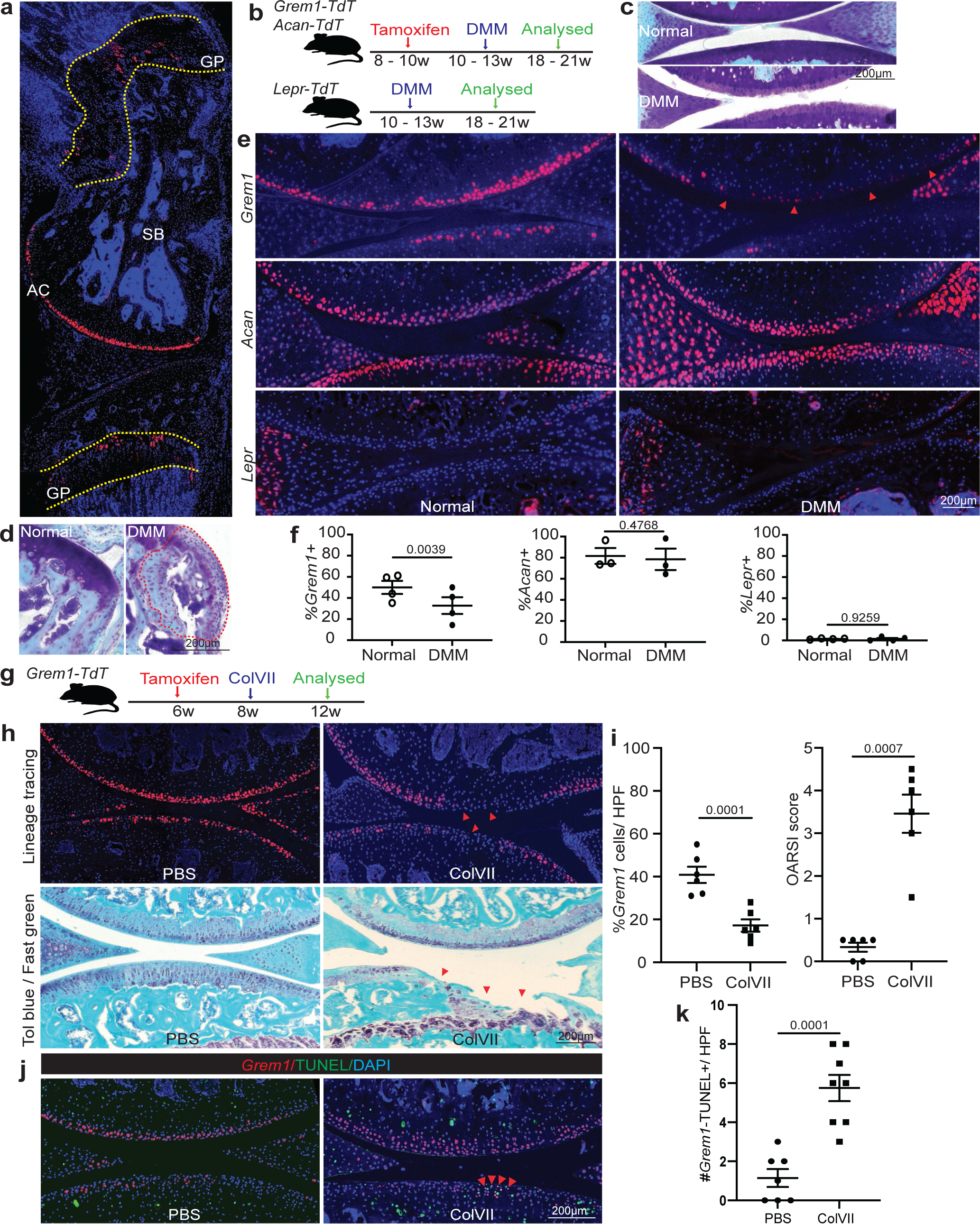
*Grem1* CP cells are depleted in OA. **(A)** Representative image of knee joint from 8 week old *Grem1-TdT* mice administered tamoxifen at 6 weeks of age showing the location of *Grem1* cells in growth plate (GP), subchondral bone (SB) and articular cartilage (AC). **(B)** Experiment schema. DMM surgery was performed on adult *Grem1-TdT*, *Acan-TdT* and *Lepr-TdT* mice and tissue harvested after 8 weeks. **(C)** Representative image of proteoglycan loss, and **(D)** osteophyte-like formation (red dotted line) stained with Toluidine blue and Fast green in DMM with paired normal for comparison. **(E)** Representative images of paired distal femur joints of *Grem1-TdT* (top), *Acan-TdT* (middle) and *Lepr-TdT* (bottom) mice showing a decrease in *Grem1*-lineage cells within the AC DMM injury site (arrows) compared to normal. **(F)** Quantification of *Grem1*-, *Acan*- and *Lepr*-lineage cells as a percentage of total chondrocytes within the DMM injury site (•) in comparison to no surgery (○) control. n=4-5 individual animals per group, Paired t test. **(G)** ColVII induced OA experiment schema. **(H)** Representative images of *Grem1-TdT* distal femur joints showing loss of *Grem1*-lineage AC cells as indicated by arrows within the injury site (top), and OA pathology induced by ColVII compared to PBS control. Sections stained with Toluidine blue and Fast green, arrows indicate superficial lesions. **(I)** Quantification of the percentage of *Grem1*-lineage cells per HPF showing significant loss of *Grem1* AC cells (left) and, unblinded histopathological assessment using OARSI grading showing significant increase in OA pathology (right) in ColVII induced OA (•) compared to PBS controls (▪). Unpaired t test. **(J)** Representative images of TUNEL staining showing increased apoptosis in articular *Grem1*-lineage cells. **(K)** Quantification of the number of *Grem1*-lineage TUNEL positive cells in ColVII induced OA (•) compared to PBS control (▪) showing a significant increase in articular *Grem1*-lineage cell death. Unpaired t test.

### *Grem1* marks a chondrogenic progenitor population in the AC with osteoblastic lineage potential

Given that OA models were characterised by loss of *Grem1*-lineage CP cells, we tested whether *Grem1*-lineage CP cells were a resident stem-progenitor cell for normal articular cartilage post-natal development and maintenance. *Grem1-TdT* and *Acan-TdT* mice were administered tamoxifen at postnatal day 4 to 6 before sacrifice with age matched *Lepr-TdT* mice, to determine the contribution of each lineage to the AC (**Figure 2A**). *Grem1*-lineage CP cells were immediately observed within the cartilaginous epiphysis and meniscus after a week, and had given rise to 39.4% of the AC. In later stages of development, we found the *Grem1*-lineage CP cells also generated osteoblasts in subchondral bone, and in one month had populated the entire joint structure, including 30.7% of the AC (**Figure 2B**). A partially overlapping distribution of AC cells was observed with *Acan* progeny (**Figure 2B**). In contrast, *Lepr*-lineage cells (Pittenger et al., 1999) were not evident within the AC but were found as perisinusoidal cells in the bone marrow, consistent with a previous report (Ding et al., 2012) (**Figure S2B**). Immunofluorescence staining at 20 weeks (**Figure 2C and S2A**) showed that *Grem1* and *Acan* cells gave rise to SOX9+ and hypertrophic COLX+ chondrocytes, as well as OCN+ osteoblasts. In contrast, *Lepr* cells only gave rise to OCN+ osteoblasts, very occasional SOX9+ chondrocytes and no COLX+ chondrocytes (**Figure 2C and S2A**).

**Figure 2.**
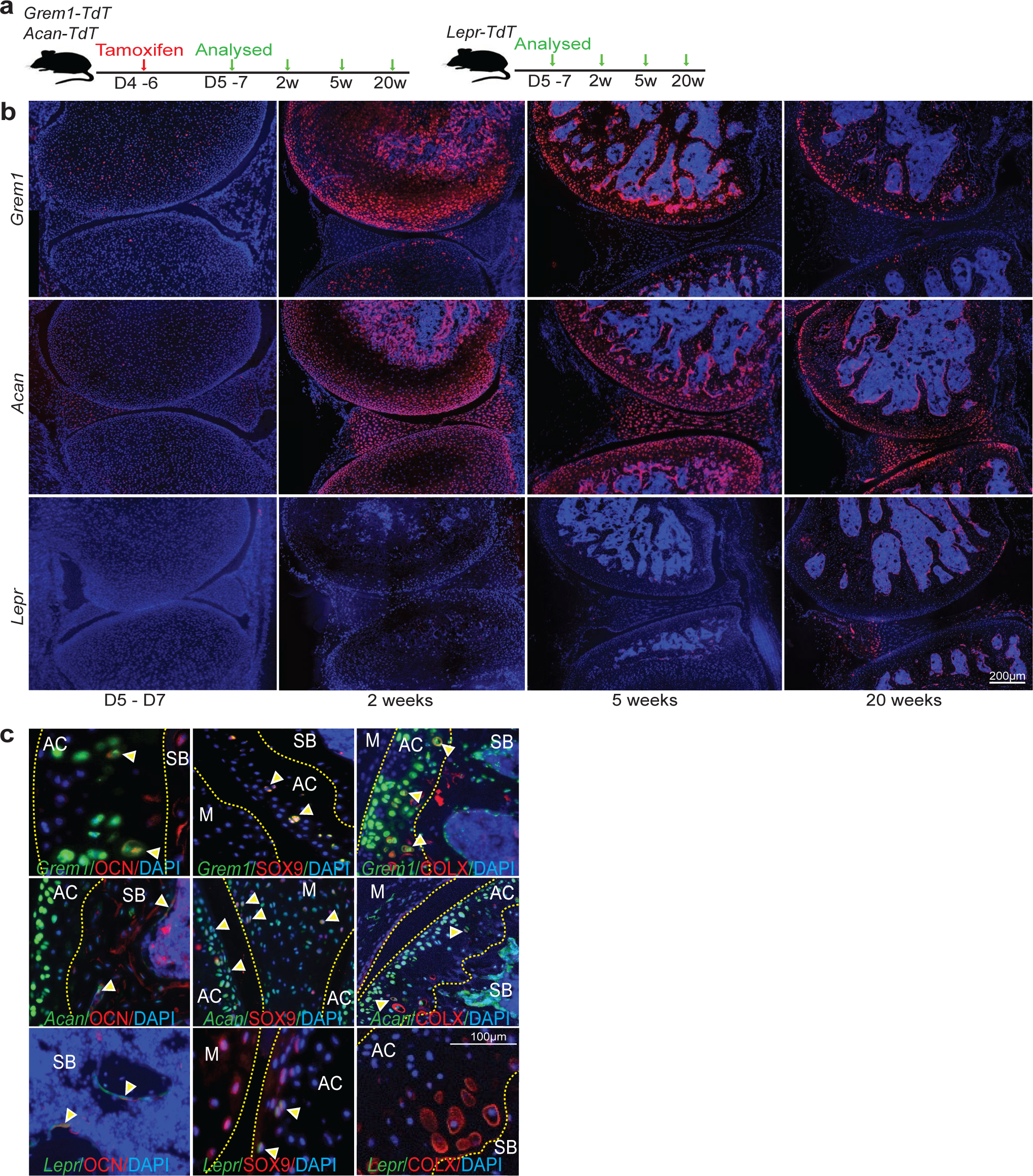
*Grem1* marks a chondrogenic progenitor population in the AC with osteoblastic lineage potential. **(A)** Experimental schema. **(B)** Representative images of AC from *Grem1-TdT* (top row) and *Acan-TdT* (middle row) mice administered tamoxifen at P4 - P6 of age, and age paired *Lepr-TdT* (bottom row) mice analysed at indicated times using fluorescence microscopy. n=5 animals per group per time point. **(C)** Histological analysis of distal femur from neonatal tamoxifen *Grem1-TdT*, *Acan-TdT* and *Lepr-TdT* pulse-chase for 20 weeks. Representative IF staining of *Grem1* (top row), *Acan* (middle row) and *Lepr* (bottom row) cells expressing OCN, SOX9 and COLX (indicated by yellow arrows). Subchondral bone (SB), articular cartilage (AC) and meniscus (M). n=3 animals per group.

### *Grem1*-lineage cells have multilineage differential potential

To further analyse the clonogenic and differentiation potential of *Grem1*-lineage CP cells in the early adult knee *ex vivo*, *Grem1-TdT* and *Acan-TdT* mice were administered tamoxifen at 6 weeks of age and knee joints harvested 2 weeks later (**Figure 3A**). *Grem1* labelled specific articular surface cells (**Figure 3B**) overlapping with, but more limited than, the total *Acan* chondrocyte population (**Figure S3B)**. These *Grem1-* and *Acan-*traced cells from the AC of the tibiofemoral joint were isolated via flow cytometry and plated at clonal density (**Figure 3C**). As *Lepr* cells were not found in the AC (**Figure S2B**), a *Lepr* comparable population was not available for this GP cartilage excluded experiment. *Grem1*-lineage articular CP clones could be serially propagated, while *Acan* clones lost serial propagation (**Figure 3C**). *Grem1* expression was also significantly higher in *Acan* clones capable of *in vitro* expansion compared to those that were not (**Figure S3A**), suggesting *Grem1* expression correlated with self-renewal *in vitro*. Of the *Grem1-* and *Acan-*lineage clones that underwent >3 passages, no significant difference in CFU-F efficiency was observed (**Figure 3D**). When subjected to multilineage differentiation, both adult *Grem1-* and *Acan-*lineage clones gave rise to osteogenic (alizarin red stain+) and chondrogenic (alcian blue stain+), but not adipogenic (oil red O stain+) progeny (**Figure 3E and S3C**), consistent with previous studies on postnatal whole bone populations (Worthley et al., 2015) (**Figure 3F**). A significantly greater percentage of *Grem1*-lineage clones were capable of osteogenic differentiation compared to *Acan*-lineage clones (100% vs 25%).

**Figure 3.**
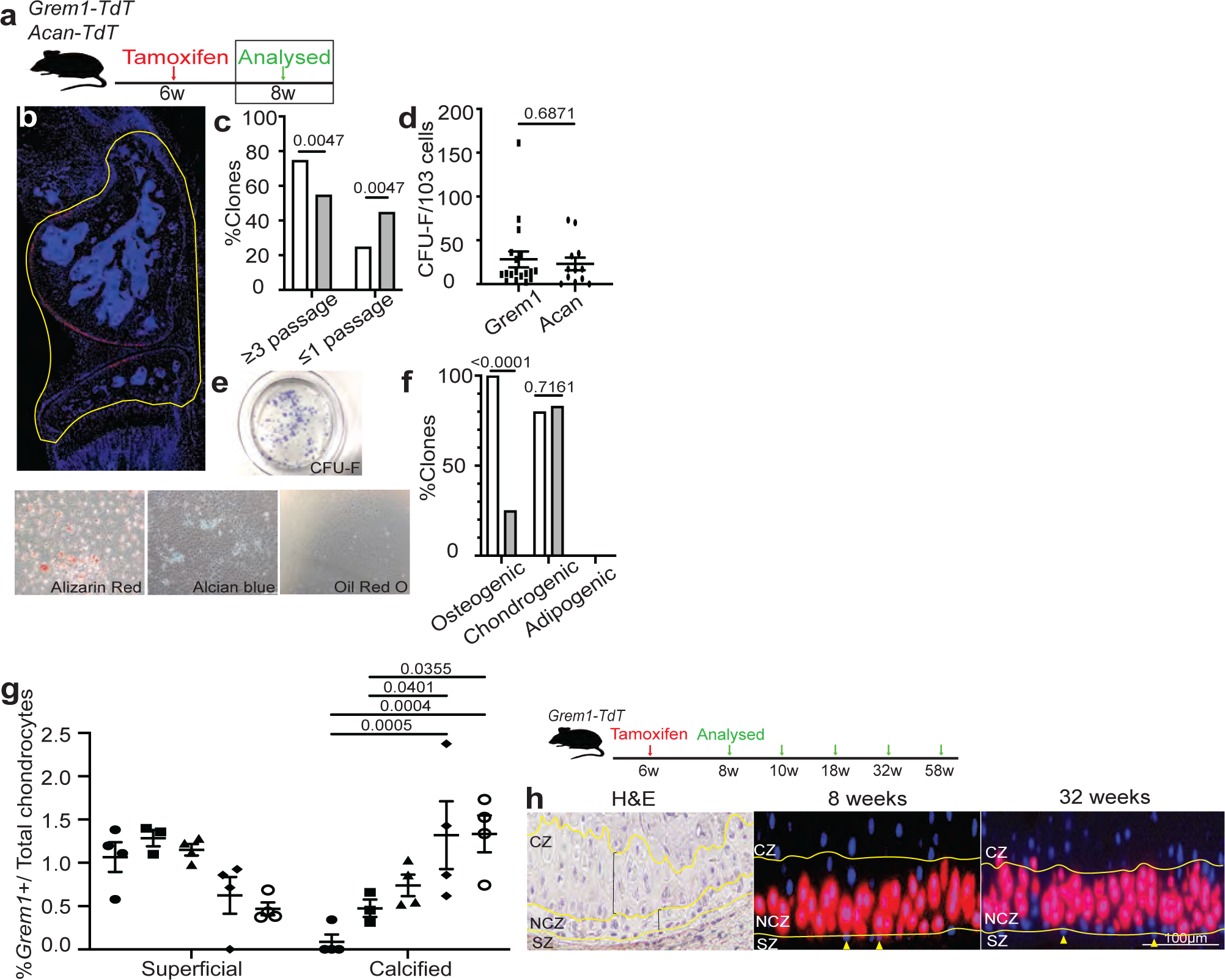
*Grem1*-lineage cells have multilineage differential potential. **(A)** Experimental schema. **(B)** Articular joints, outlined in yellow, were used to isolate red cells for *in vitro* assays using flow cytometry. Pooled cells from n=3 animals per cell population were seeded at clonal density with 22 – 24 clones per cell population assayed. **(C)** Percentage of *Grem1*-lineage (white) clones able to undergo expansion compared to *Acan*-lineage (grey) clones. **(D)** Number of CFU-F formed per clone, each data point represents an individual clone. **(E)** Representative images of *Grem1*-lineage cells stained for CFU-F or differentiation markers Alizarin Red (osteo), Alcian blue (chondro) and Oil Red O (adipo). **(F)** Number of clones that had undergone osteogenic, chondrogenic, and adipogenic differentiation quantified as a percentage of the total number of clones isolated. **(G)** Experimental Schema (right). Tamoxifen was administered to *Grem1-TdT* mice at 6 weeks of age and pulse-chased for 12 months. Quantification of the total number of *Grem1*-lineage cells within the superficial and calcified zones as a percentage of the total number of chondrocytes within the AC at 8 weeks (•), 10 weeks (▪), 18 weeks (▴), 32 weeks (♦) and 58 weeks (○) of age showing a significant increase in *Grem1*-lineage AC cell contribution to the calcified zone with age (left). n=3-4 mice per time point, each data point represents an individual animal. **(H)** Representative images of adult articular joint showing H&E of zonal organisation of chondrocytes in the superficial zone (SZ), non-calcified zone (NCZ) and calcified zone (CZ) and *Grem1*-lineage AC cells moving towards the CZ from 8 weeks (middle) to 32 weeks (right) of age. Superficial chondrocytes indicated with yellow arrows. **c**, **f,** Fisher’s exact test, **d,** unpaired t test, **g**,Two-way Anova Tukey’s test.

Normal AC development and maintenance requires both interstitial and appositional growth of the AC (Li et al., 2017). Using long-term cell fate tracing of adult *Grem1*-lineage cells *in vivo*, we observed these cells contributing to progenitor populations in the superficial zone of the AC and with a significant increase in the percentage of *Grem1*-lineage cells within the calcified zone of the AC with age (**Figure 3G and 3H**).

### *Grem1*-lineage articular CP cells are lost with age

To measure the longevity of *Grem1* lineage cells within the tibiofemoral joint, 6-week-old *Grem1-TdT* mice were administered tamoxifen in early adulthood and analysed during aging (**Figure 4A**). Quantification of fluorescence images showed a significant decrease in *Grem1*-lineage articular CP cells from young to aged adult mice (**Figure 4B and 4C**). This is consistent with other studies looking at the presence and regenerative capacity of skeletal stem cells in aged animals (Ambrosi et al., 2021; Murphy et al., 2020). The reduced regenerative capacity of the AC with age is due, at least in part, to reduced *Grem1*-lineage articular CP cells and chondrocyte proliferation (**Figure S4A and S4B**). We next examined whether *Grem1*-lineage CP cells persist in the aging knee by administration of tamoxifen to *Grem1-TdT* mice at 29 weeks-of-age, in comparison to postnatal (week 1) or early adulthood (6 weeks), and visualisation of lineage traced cells two weeks later (**Figure 4D**). *Grem1*-lineage cell number decreased significantly with increasing adult age, with only <0.4% of total articular surface cells being *Grem1*-lineage cells in early middle-age (**Figure 4E and 4F**). At this stage, *Grem1*-lineage cells were mainly observed in the subchondral marrow space and occasionally in the retained but no longer actively growing GP (**Figure S4C**).

**Figure 4.**
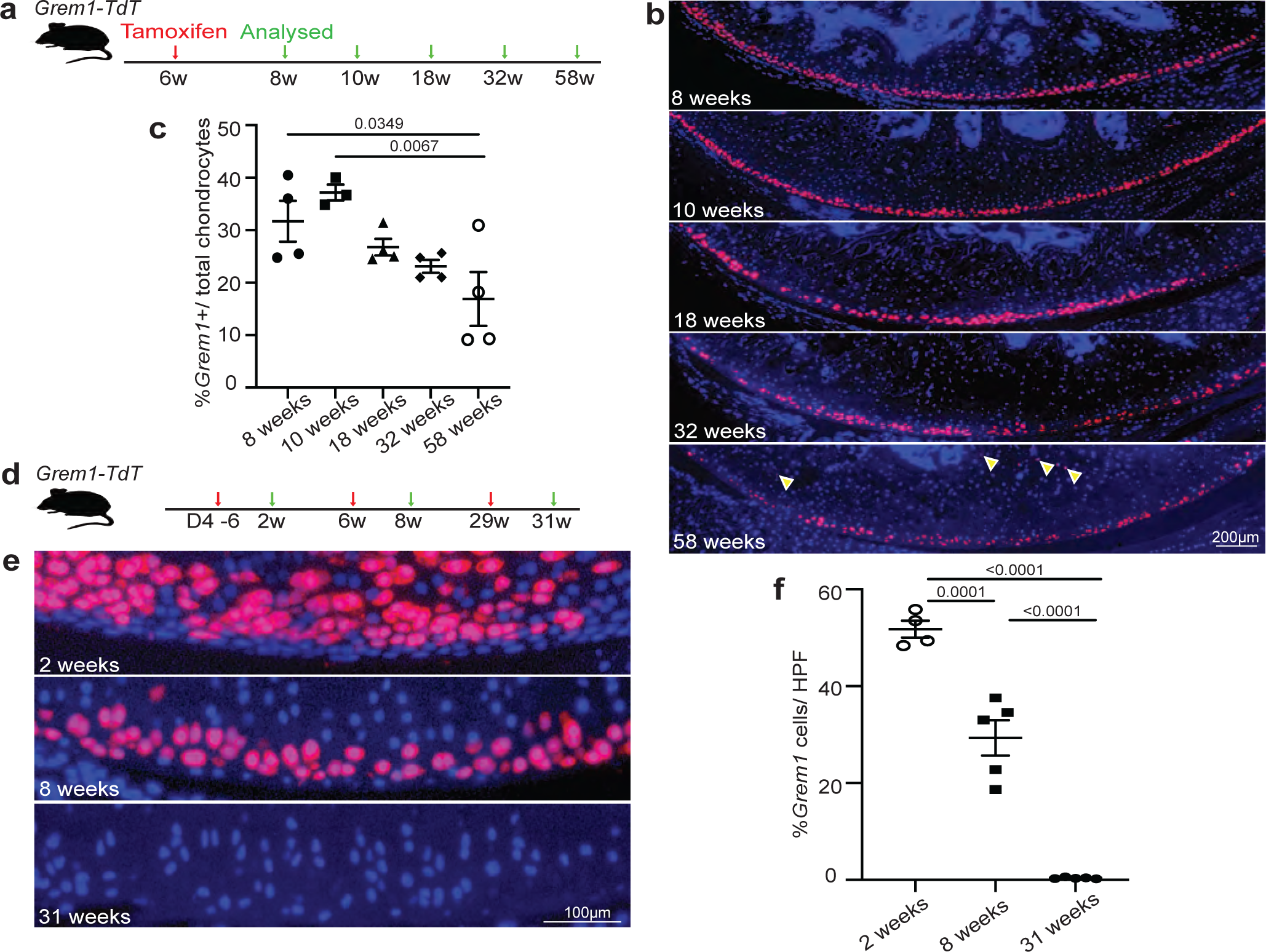
*Grem-*1lineage articular CP cells are lost with age. **(A)** Experimental schema. **(B)** Representative images of the articular knee joint collected at indicated times. An increased number of *Grem1*-lineage cells were observed in the subchondral bone area indicated by yellow arrows at 58 weeks of age. **(C)** Quantification of the total number of *Grem1*-lineage articular chondrocytes as a percentage of the total number of articular chondrocytes showing a significant decrease in *Grem1*-lineage articular chondrocytes with age. n=3-4 animals per time point, data point represents an individual animal analysed. One-way Anova Tukey’s test. **(D)** Experiment schema. **(E)** Representative images of *Grem1*-lineage articular cells within the articular joints of mice at different postnatal stages, n=4-5 animals per time point. **(F)** Quantification of the total number of *Grem1*-lineage articular chondrocytes as a percentage of the total cells per high power field (HPF) in different postnatal stages showing a significant loss of *Grem1*-lineage AC cells in aged articular knee joints. Data point represents an individual animal analysed. One-way Anova Tukey’s test.

### Targeted ablation of *Grem1* cells causes OA

We have previously examined the role of *Grem1*-lineage cells in postnatal skeletogenesis using a diphtheria toxin ablation model (*Grem1*-*creERT;R26-LSL-ZsGreen;R26-LSL-DTA*), in which the *Grem1*-lineage was incompletely ablated(Worthley et al., 2015). Nevertheless, post-natal ablation of *Grem1*-lineage cells led to significantly reduced femoral bone volume by microCT after two weeks, with reduced trabecular bone as quantified by histology (Worthley et al., 2015). To increase the efficiency of *Grem1*-lineage ablation and investigate the consequences of adult *Grem1* CP cell loss in the AC, we first generated a new *Grem1-DTR-TdTomato* knock in mouse model (*Grem1-DTR-Td)* in which *Grem1* cells express the TdTomato reporter and Diphtheria toxin receptor (DTR), thus making them susceptible to diphtheria toxin (DT) ablation. These mice were administered 2 doses of DT intra-articularly at 5 to 6 weeks of age and sacrificed 3 days later (**Figure 5A and S5A**). This induced a significant reduction in *Grem1* CP cell number in the AC but not GP (**Figure 5B, 5C and S5B**), increased COLX+ articular chondrocytes and increased blinded OA pathology scoring in DT treated *Grem1-DTR-TdTomato* mice compared to age-matched control groups (**Figure 5D and 5E**). This increase in OA score with *Grem1*-cell ablation was predominantly due to cartilage hypertrophy, proteoglycan loss and structural damage, with a lesser contribution from meniscus pathology and subchondral vascular invasion and zero scoring from subchondral bone sclerosis or osteophyte changes (**Figure 5F-5J**). To confirm the role for adult *Grem1* CP cells in OA we utilised a second transgenic mouse model of targeted cell ablation generated by mating *Grem1-TdT* mice to *Rosa-iDTR* mice to create *Grem1-TdTomato-iDTR* mice (*Grem1-creERT;DTR)*. DTR expression on *Grem1*-lineage cells was induced by tamoxifen administration at 4 weeks of age, followed by local ablation of *Grem1*-lineage AC cells by intra-articular injection of DT at 8-weeks-old for 2 weeks, before animals were sacrificed at 26 weeks (**Figure S5C**). Analysis of OA pathology and quantification of *Grem1*-lineage articular surface CP cells showed a significant loss in *Grem1*-lineage AC cells concomitant with worsened OA pathology, including a reduction in AC thickness (**Figure S5D and S5E**). Together our data confirmed that *Grem1*-lineage CP cells are normal progenitor cells that are lost in aging and in mechanical trauma that causes OA and their depletion directly results in OA.

**Figure 5.**
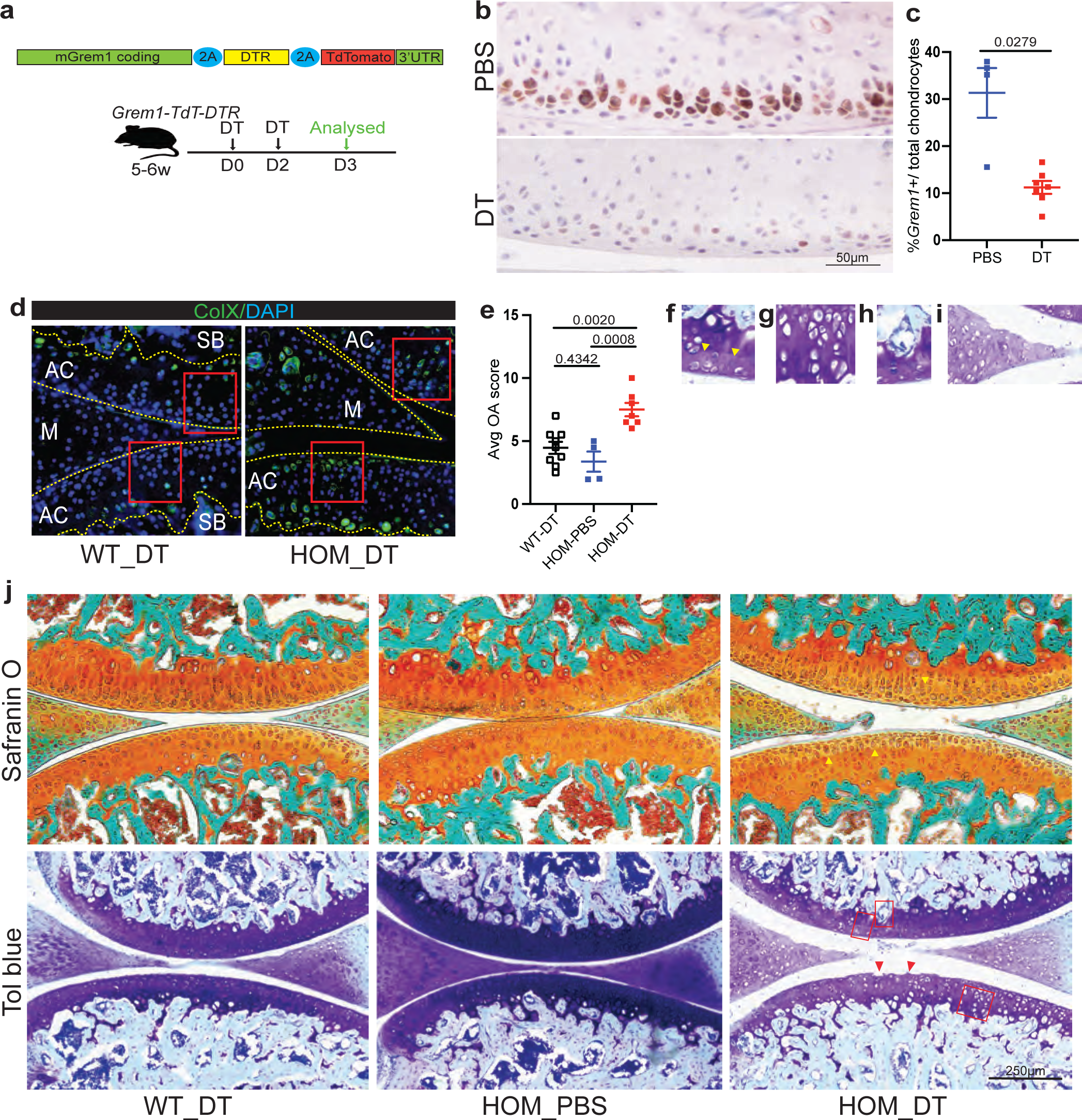
Targeted ablation of *Grem1* cells causes OA. **(A)** A new knock-in *Grem1* ablation mouse (*Grem1-Td-DTR*) was generated and used to investigate the role of the *Grem1* in the AC. Schematic showing the *Grem1-Td-DTR* knock-in construct (top) and experimental outline involving targeted ablation of *Grem1*-expressing cells in the AC achieved via intra-articular injection of DT into adult *Grem1-Td-DTR* mice (bottom). **(B)** Representative images of early adult mice articular joints treated with PBS or DT stained with anti-RFP. **(C)** Quantification of *Grem1* AC cells as a percentage of the total number of AC chondrocytes in PBS (blue) or DT treated (red) *Grem1-Td-DTR* mice, showing a significant decrease with DT indicative of successful ablation of *Grem1* cells. Welch’s t test. **(D)** Representative images of ColX staining of Grem1-Td-DTR joints from WT and HOM DT-treated animals. Boxed regions indicate loss of ColX chondrocytes following ablation of Grem1-expressing articular cells in HOM DT treated mice. **(E)** Blinded scoring of average OA score following 3 days of targeted ablation of *Grem1* AC cells in homozygous *Grem1-Td-DTR* mice showed a significant increase in OA pathology compared to controls. Each data point represents an individual animal analysed, n=4-9 per group. One-way Anova Tukey’s test. **(F – I)** Representative image of DT treated articular joint of *Grem1-Td-DTR* mice stained with Tol blue and fast green showing pathological changes commonly associated with OA **F**, loss of proteoglycan in the AC indicated by yellow arrows, **G**, hypertrophic chondrocytes, **G**, SB invasion, and **I**, meniscus pathology. **(J)** Representative images of *Grem1-Td-DTR* joints from WT treated with DT, and HOM treated with PBS or DT, stained with Safranin O (top) or Tol Blue (bottom) and fast green showing proteoglycan loss as indicated by decreased staining intensity with the orange (Safranin O, yellow arrowheads) and purple (Tol Blue) stains in HOM DT-treated animals. HOM DT-treated also showed signs of AC damage indicated by red arrows. Red boxes represent OA pathology indicated in F – I.

While we showed that loss of *Grem1*-lineage CP cells is an early event in OA (**Figure 1**) and in turn also causes OA (**Figure 5**, **S5D-E**), we considered the possibility that OA may also be caused by the death or degeneration of *Grem1-*lineage, mature Acan-expressing chondrocytes. Previous studies noted that ablation of chondrocytes marked by *Acan* or *Prg4* results in mild OA that resolves over time, or no OA (Masson et al., 2019; Zhang et al., 2016). To further discriminate the role of *Acan+* chondrocytes in comparison to *Grem1*-lineage CPs in OA, we compared the OA phenotype caused by ablation of each cell population using similar genetic models and induction regimens. In *Acan-creERT;DTR* mice, immunofluorescent staining for ACAN showed strong expression in the more mature chondrocytes 2-3 cells below the articular surface, with limited expression in cells at the articular surface (**Figure S5F**). Fortuitously, DT injection into the knee joints of *Acan-creERT;DTR* mice caused ablation of the ACAN-bright chondrocyte population, while the leaving the articular surface layer relatively intact, and thereby enabled us to assess ablation of the ACAN-bright chondrocytic cells. DT treatment of *Acan-creERT;DTR* mice caused a significant decrease in ACAN-expressing cells, but only a comparatively mild numerical increase in OA pathology scoring at 26 weeks that was not statistically significantly different to saline treated controls (**Figure S5F-G**). The overall average OA score in *Acan*-lineage ablated mice was 0.5 (+/- 0.25, st.dev.) in comparison to 1.75 (+/- 0.68) in the *Grem1-TdT-DTR* mice treated with DT (**Figure S5D-G**). These data are consistent with previous *Acan-* and *Prg4-* chondrocyte ablation experiments reported to generate mild or insignificant OA (Masson et al., 2019; Zhang et al., 2016). Given these differences, we note that OA is not just a disease of mature chondrocyte loss, but is also likely also a failure of local progenitor regeneration.

We next used flow cytometry to isolate *Grem1*-lineage cells from early adult mice and implanted them into recipient mice that had undergone DMM surgery via intra-articular injection. This initial effort to implant *Grem1*-lineage cells was hindered by inefficient homing to and/or survival of the cells in the AC (**Figure S5H-I).** Thus, we subsequently studied how to preserve this important population through normal aging (**Figure 2**) or injury associated (**Figure 1**) attrition that could be key for OA prevention.

### *Grem1*-lineage single cell transcriptomics revealed a distinct population of articular chondrocytes

We applied scRNAseq analysis to characterise FACS-isolated early adult *Grem1*-lineage CP cells from the epiphysis proximal to the GP and compared to *Grem1*-lineage cells within the GP, as well as age-matched skeletal cells defined by the *Lepr* lineage (**Figure S6A and S6B**). Unsupervised clustering of our pooled scRNAseq data revealed 6 distinct cell clusters: five *Grem1*-lineage (AC1-2, GP1-3) and one *Lepr* cluster (**Figure 6A, 6B and S6C-S6E**). Though there were substantial similarities in transcript expression in the articular surface and GP cells, clusters within the *Grem1*-lineage cells in the articular surface could also be separated by differential transcript expression from those in the GP (**Figure 6A, 6B, S4F and Table S1**). Of the *Grem1*-lineage cells, AC cluster 1(AC1) showed differential expression of a cytoprotective marker, *Clusterin* (Connor et al., 2001), and early chondrogenic differentiation marker, *Integral membrane protein 2A* (Van den Plas and Merregaert, 2004); AC cluster 2 (AC2) the hypertrophic chondrocyte marker *Col10a1* (Kielty et al., 1985) and early chondrocyte marker *Sox9*(Lefebvre et al., 2019); GP cluster 1 (GP1) and 2 (GP2) the cartilage angiostatic factor *Chrondromodulin-1* (*Cnmd*), known for its role in endochondral ossification and bone formation (Shukunami and Hiraki, 2001; Zhu et al., 2019), *melanoma inhibitory factor* (*Mia*) (Bosserhoff and Buettner, 2002) and *Matn3* which encodes an ECM protein that regulates cartilage development and homeostasis (Jayasuriya et al., 2012) while GP1 also expressed cartilage differentiation marker *Serpina1* (Boeuf et al., 2008); GP cluster 3 (GP3) expressed cartilage degeneration marker *Mmp13* (Burrage et al., 2006) (**Figure 6B, S6E and Table S1**). Consistent with previous studies, this included enrichment of the AC marker, *Lubricin* (*Prg4*), in AC compared to GP *Grem1*-lineage cells (Kozhemyakina et al., 2015) (**Figure S6G**). Immunofluorescence staining of tissue samples from two week lineage traced *Grem1-TdT* mice showed overlap between articular *Greml*-lineage CP and PRG4 expressing cells (12.6% of the articular cartilage double positive), but the *Greml*-lineage cells were largely distinct from COL2-expressing chondrocytes in the calcified zone (<0.1% articular cartilage cells double positive for *Grem1*-lineage and COL2, **Figure 6C**). Most *Grem1*-lineage cells separately prepared from whole bone expressed the defining marker of mSSC, CD200(Chan et al., 2015) (**Figure S6H**), but only 19% of *Grem1*-expressing cells from the articular surface had a transcriptomic profile consistent with the mSSC immunophenotype (**Figure S6I and S6J**). This indicates that *Grem1*-lineage CP in the articular surface and mSSC in the GP partially overlap, but the articular surface *Grem1*-lineage population is distinct.

**Figure 6.**
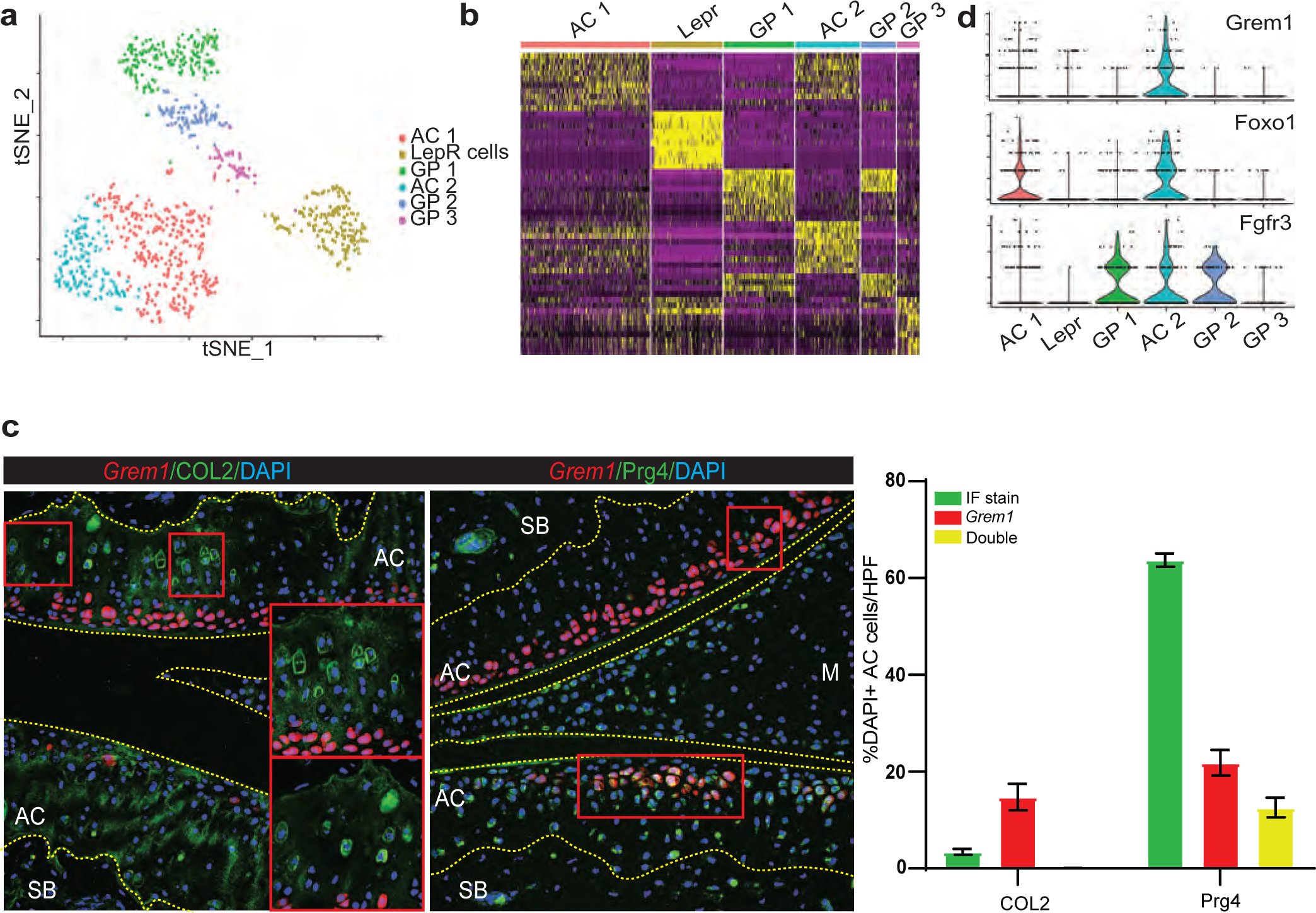
*Grem1*-lineage single cell transcriptomics revealed a distinct population of articular chondrocytes. **(A)** Single cell RNA (scRNA) sequencing data showed distinct clusters of cells isolated from the AC and GP of *Grem1-TdT* mice compared to *Lepr*-lineage cells isolated from the *Lepr-TdT* mice. **(B)** Heat map depicting unsupervised clustering of top 10 differentially expressed transcripts between the different clusters in (**A)**. **(C)** Representative images of IF staining of *Grem1-TdT* joints induced at 6 weeks and collected at 8 weeks old. Red squares highlight regions of interest with positive staining. COL2 staining of the AC showed little overlap between COL2 expressing chondrocytes and *Grem1*-lineage cells (yellow). PRG4 staining of the AC however, showed a larger population of *Grem1-*lineage cells expressing PRG4 (yellow). Quantification of total COL2, PRG4 and *Grem1-*lineage cells represented as a percentage of total AC chondrocytes. Only cells that were positive for DAPI were counted to ensure that only live cells were quantified. The percentage of *Grem1*-lineage cells that express COL2 was <0.1%, indicative of 2 distinct populations of cells. 64% of AC chondrocytes expressed PRG4 and about half of the *Grem1*-lineage cells also express PRG4. Quantification performed using n=5 animals per group. **(D)** *Grem1*-lineage AC cells co-expressed genes important for AC function (*Foxo1*) and receptor (*Fgfr3*) for FGF18 treatment. Chi-Square correlation analysis confirmed co-expression of *Foxo1* and *Fgfr3* in *Grem1* expressing cells (p=0.00113).

### Target for OA therapy identified through single cell transcriptomics

Next, we investigated gene expression associated with cartilage development and regeneration across the scRNAseq data set. *Forkhead box protein-o* (*Foxo)* genes have important roles in apoptosis, cell-cycle progression, resistance to oxidative-stress and maintenance of stem cell regenerative potential (Miyamoto et al., 2007; Zhang et al., 2011). We observed that *Foxo1* expression was highly correlated with *Grem1* expression (p<0.0001) in articular CP cells in our scRNAseq data (Matsuzaki et al., 2018) (**Figure 6D**), with *Foxo1* expression also significantly increased in the AC compared to GP *Grem1*-lineage populations (**Figure S7A**). At the protein level, the majority of FOXO1 expressing cells in the adult AC were *Grem1*-lineage cells, while there were very few FOXO1 expressing *Grem1*-lineage cells observed in the GP (**Figure S7B**). To determine whether *Foxo1* expression was essential for survival of *Grem1* articular CP cells and prevention of OA, *Grem1-TdT* mice were crossed to *Foxo1^fl/fl^* mice, to produce *Grem1-TdT-Foxo1* conditional knock-out mice. *Grem1-TdT-Foxo1* mice were administered tamoxifen at 6 weeks of age to induce *Foxo1* deletion in *Grem1*-lineage cells and sacrificed at 26 weeks (**Figure 7A**). Significantly fewer *Grem1*-lineage AC but not GP cells were observed in *Grem1-TdT-Foxo1* mice, in comparison to age-matched *Grem1-TdT* controls, and the resulting increased OA pathology resembled *Grem1-creERT;DTR* DT ablation (**Figure 7B-7D and S7C**). This suggested that *Foxo1* was important to maintain adult *Grem1*-lineage articular CP cells and, by extension, AC integrity. Interestingly, *Grem1-TdT-Foxo1* mice administered tamoxifen in the early neonatal period (Day 4-6) developed drastic *Grem1*-lineage articular CP cell loss, more severe OA pathology and decreased AC thickness, in comparison to age-matched *Grem1-TdT* controls and *Grem1-TdT-Foxo1* animals induced with tamoxifen in early adulthood (**Figure S7D and S7E**). This indicates the crucial role for *Foxo1* in AC development of the *Grem1*-lineage CP cell population, as well as in adult AC maintenance.

**Figure 7.**
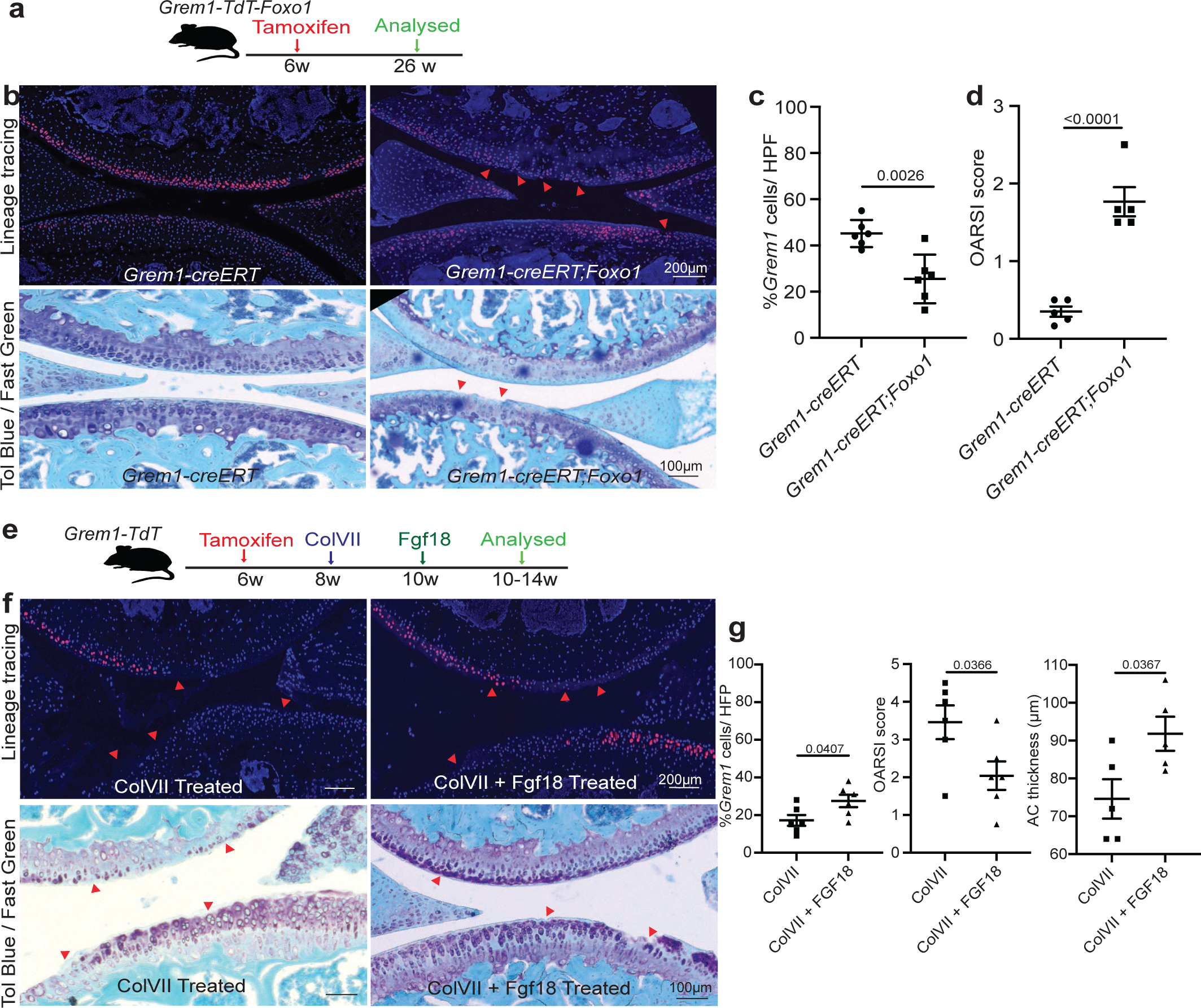
OA can be caused by loss of *Foxo1* in *Grem1*-lineage CP cells and partially rescued by FGF18 treatment. **(A)** Experiment schema. **(B)** Representative images of fluorescent *Grem1* lineage tracing in the articular joint with or without *Foxo1* deletion (top) with OA lesions highlighted using red arrows, and Toluidine blue and Fast green stain (bottom) with arrows indicating cartilage lesions and chondrocyte disorganisation. n=5 animals per group. **(C)** Percentage of *Grem1*-lineage articular cells per HFP in control *Grem1-TdT* mice (•) compared to *Grem1-TdT-Foxo1* mice (▪) showed a significant decrease in *Grem1*-lineage cells with loss of *Foxo1*. **(D)** Unblinded histopathological OARSI scoring showed a significant increase in OA score in *Grem1-TdT-Foxo1* mice (▪) compared to *Grem1-TdT* mice (•). Each data point represents an individual animal analysed. **(E)** Experimental schema. **(F)** Representative images of joints from *Grem1*-lineage mice with ColVII induced OA with or without FGF18 treatment with arrows indicating injury site (fluorescence, top), toluidine blue and fast green stained showing proteoglycan loss and lesions indicated by arrows (bottom). **(G)** Quantification of the percentage of *Grem1*-lineage cells showed a significant increase with FGF18 treatment (left) which resulted in a delay in OA progression (middle) and rescued AC thickness (right). n=5-6 animals per group, each data point represents an individual animal analysed. All statistical analyses Unpaired t test.

To discover molecular targets in *Grem1*-lineage articular CP cells for OA therapy we focussed on fibroblast growth factor (FGF) signalling, given the role of this pathway in regulating chondrogenesis (Ornitz and Marie, 2015). *Fibroblast growth factor receptor 3* (*Fgfr3)* was expressed both in *Grem1*-lineage GP and articular CP populations. Aware of the important role for fibroblast growth factor 18 (FGF18) as a key agonist of FGFR3 as an experimental treatment for OA (Liu et al., 2007), we investigated the impact of exogenous FGF18 on *Grem1*-lineage AC stem cells *in vivo*. Adult *Grem1-TdT* mice induced with tamoxifen were injected intra-articularly with 0.5μg FGF18 or PBS twice weekly for 2 weeks and concurrently administered EdU to label newly proliferating cells (**Figure S7F**). FGF18 treatment resulted in a significant increase in both the number of EdU+ *Grem1*-lineage articular CP cells and AC thickness, suggesting FGF18 induces AC chondrogenesis in the adult knee via increased *Grem1*-lineage articular CP cell proliferation (**Figure S7G and S7H**). Notably, very few EdU+ cells were found in the hypertrophic chondrocyte zone, suggesting that the neo-cartilage arises from the articular CP population. To determine whether FGF18 can also rescue OA pathology, we used ColVII to induce OA and then intra-articularly injected *Grem1-TdT* mice with 0.5μg FGF18 or PBS twice weekly for 2 weeks. FGF18 injection resulted in a significant increase in *Grem1*-lineage CP cells and AC thickness in treated joints compared to PBS controls and a significant reduction in OA pathology (**Figure 7E-7G**). This suggested that exogenous FGF18 alleviates OA through increased *Grem1*-lineage CP cell proliferation.

## DISCUSSION

OA, like other degenerative diseases, can be viewed as the dysfunction of mature cartilage cells or the result of an imbalance in stem-progenitor cell repair and renewal. Previous studies have shown functional cartilage cell dysfunction/death results in OA, here we show that ablation of a restricted progenitor population can cause OA without disturbing the broader population of differentiated chondrocytes.

While stem cell therapy for OA using traditional MSC populations has shown promise in reducing pain and stiffness, it does not effectively repair normal AC (Feczko et al., 2003; Hangody and Fules, 2003). Several research teams, including our own, have recently described SSCs with lineage potential restricted to cartilage, bone and stroma that are promising candidates for joint disease treatment (Chan et al., 2015; Murphy et al., 2020; Worthley et al., 2015). In this study, we show that *Grem1*-lineage CP cells contribute to neonatal generation of the AC and are pivotal to AC maintenance in adulthood. *In vitro*, the articular *Grem1*-lineage cells display clonogenicity and multi-potent differentiation properties associated with stemness, however a complete assessment of their potential stem cell properties and lineage hierarchy using *in vivo* transplantation studies (Chan et al., 2015; Chan et al., 2018) remains to be undertaken.

The position of adult *Grem1*-lineage traced cells on the superficial surface of the AC, and subsequently in deeper layer chondrocytes and subchondral bone, is consistent with previous descriptions of chondrocytic label retaining progenitors and the *Prg4* (encoding lubricin) lineage where tracing began from birth or in juvenile mice, not adults as investigated here (Kozhemyakina et al., 2015; Li et al., 2017). A key difference between the *Prg4*- and *Grem1*-lineages in the tibio-femoral joint is that unlike *Prg4*-, *Grem1*- also gives rise to cells in the growth plate (Kozhemyakina et al., 2015). Although the AC and GP are both broadly constituted by chondrocytes, these structures form quite differently and have differing roles in skeletogenesis (Candela et al., 2014; Decker, 2017), with the AC being the primary site of OA pathology. Our focus has been on the articular *Grem1*-lineage population and superficial chondrocytes as this is the site of early OA-like changes. We cannot discount, however, that the GP *Grem1*-lineage CP cells may also contribute to the articular phenotypes observed and have not examined whether *Grem1*-lineage subchondral bone populations arise from GP or AC progenitors or both. We show that some, but not all, acutely labelled *Grem1*-lineage articular cells express *Prg4*/PRG4 and are distinct from the COL2-expressing chondrocytes of the calcified zone. Other key differences between the *Grem1*-lineage and that of the broader chondrocytic population marked by *Acan* include the extent and location of labelled lineage cells (**Figure 2B and S3B**), clonogenicity and differential potential *ex vivo* (**Figure 3A-3F**) and response to injury (**Figure 1B-F**), such that *Grem1*- but not *Acan*-lineage cells were lost in the early OA-like phenotype induced by DMM surgery. Ablation of *Grem1*-lineage CP cells in the knee joint causes OA, with a much milder to no phenotype observed using identical induction protocols and similar transgenic mouse models to ablate *Acan*-lineage cells (**Figure S5D-G**). Likewise, as FOXO1 expression is restricted to the superficial chondrocytes of the AC (**Figure S7B)**, genetic deletion of *Foxo1* in *Grem1*-lineage cells effectively depletes FOXO1 expression in the articular surface, rather than deeper chondrocytic layers and results in OA (**Figure 7A-D**). This suggests the *Grem1*-lineage is a specific subset of the total chondrocyte population that may be lost early in the disease process. We acknowledge that OA can be caused by dysfunction of mature cartilage cells, but here also highlight the important function of a specific chondrocyte progenitor population marked by *Grem1*. This is supported by phenotypic differences resulting from adult *Prg4* and *Grem1*-lineage ablation. Loss of both *Prg4*- and *Grem1*-lineage populations resulted in decreased superficial chondrocytes, however cartilage deterioration and OA histological features only occurred following loss of *Grem1*-, and not *Prg4*-, lineages (Zhang et al., 2016) (**Figure 5)**.

Similar to mSSC cells (Murphy et al., 2020), *Grem1*-lineage articular CP cells were depleted with age (**Figure 4**). Using two mouse models of OA, we found that *Grem1*-lineage CP cells in the knee joint were lost in disease (**Figure 1**). For the first time, we also report that functional ablation of *Grem1*-lineage articular CP cells, or genetic deletion of *Foxo1* in *Grem1*-lineage cells, in early adult mice led to significant and rapid OA (**Figure 5,7**). As efforts to reintroduce *Grem1* articular CP cells would be a natural first step to therapeutics, our initial efforts were not successful (**Figure S5F-G**) and future efforts using a bio-scaffold with factors to stimulate enhanced chondrogenesis are warranted. scRNAseq analysis identified separate populations of *Grem1*-lineage cells within the AC and GP structures, that provided an insight to the restricted commitment of each population of cells. FGF18 treatment induced proliferation of *Grem1*-lineage articular CP cells and reduced OA pathology, presenting a strong candidate for further clinical trials of OA prevention and therapy for early disease.

We propose that the initial stage of OA can be predisposed by inadequate reserves of, or injury to, articular CP cells (accompanied by surface proteoglycan loss and fibrillation), followed by the apoptotic death of further CP cells, causing the inability to regenerate articular cartilage. This study reframes OA as a degenerative disease that can result from CP cell loss and provides a new focus for OA therapy.

## ACKNOWLEDGEMENTS

We acknowledge the facilities and the scientific and technical assistance of the South Australian Genome Editing (SAGE) Facility, the University of Adelaide, and the South Australian Health and Medical Research Institute. SAGE is supported by Phenomics Australia. Phenomics Australia is supported by the Australian Government through the National Collaborative Research Infrastructure Strategy (NCRIS) program. Flow cytometry analysis and/or cell sorting was performed at the Adelaide Health and BioMedical Precinct Cytometry Facility. The Facility is generously supported by the Detmold Hoopman Group, Australian Cancer Research Foundation and Australian Government through the Zero Childhood Cancer Program. We also acknowledge the facilities and scientific and technical assistance of the JP Sulzberger Columbia Genome Center and the Confocal and Specialized Microscopy Shared Resource of the Herbert Irving Comprehensive Cancer Center at Columbia University, USA. This research was funded in part through the NIH/NCI Cancer Center Support Grant P30CA013696 and used the Genomics and High Throughput Screening Shared Resource. This publication was supported by the National Center for Advancing Translational Sciences, National Institutes of Health, through Grant Number UL1TR001873. The content is solely the responsibility of the authors and does not necessarily represent the official views of the NIH.

This study was supported by grants from the National Health and Medical Research Council (APP1099283 to D.L.W.); Cancer Council SA Beat Cancer Project on behalf of its donors and the State Government of South Australia through the Department of Health (MCF0418 to S.L.W., D.L.W.); Endeavour Research Fellowship from the Australian Government, Department of Education and Training (ERF_RDDH_179965 to J.N.); National Institute of Health (R01 to S.M.)

## AUTHOR CONTRIBUTIONS

J.N., T.J., D.L.W., S.M. conceived and designed the study. J.N. and T.J. performed most of the experiments. C.L. developed methodology and performed blinded OA scoring. T.W. analyzed scRNAseq dataset. Y.M., L.V., M.I., T.L., N.S., J.G., H.K. assisted with animal husbandry, tissue collection. T.C.W., D.H., D.M., S.G. provided material or technical support, assisted with data analysis.

D.L.W., S.L.W. and S.M. supervised the project and procured funding. J.N., S.L.W., T.J., D.L.W., and S.M. wrote the manuscript. All authors contributed substantially to the discussion of content for the article, reviewed and/or edited the manuscript before submission.

