## Supplementary material for "Loss of *Grem1*-articular cartilage progenitor cells causes osteoarthritis": Supp Figures and Table

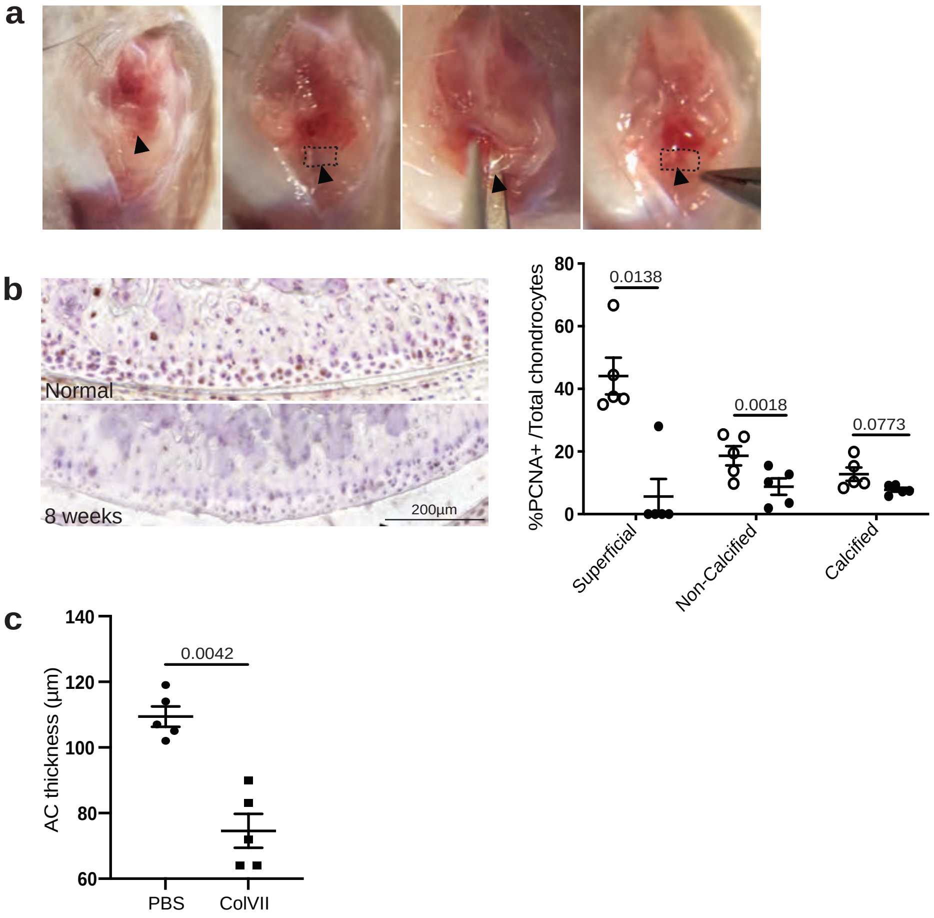


**Figure S1. Related to Figure 1**

**(A)** The destabilisation of the medial meniscus model (DMM) generates OA pathology through altered joint biomechanics, following surgical transection of the medial meniscus. Sequential images showing the transection of the medial meniscus within the knee joint (dotted box) to produce the DMM-induced OA pathology in mice.

**(B)** Representative images of PCNA immunohistochemistry in knee joints 8 weeks after DMM surgery (left) in comparison to paired, no surgery contralateral knee (normal). PCNA quantification shows a significant decrease in proliferating cells in both the superficial and non-calcified zones of the AC following DMM surgery (●) compared to no surgery (**O**), indicating no repair response 8 weeks post-surgery (right). n=5 animals per group, each data point represents an individual animal analysed. Paired t test.

**(C)** Quantification of AC thickness in the ColVII induced OA model showing a significant decrease in cartilage thickness between the control (●) and ColVII treated (■) knees. n=5 animals per group, each data point represents an individual animal analysed. Unpaired t test.

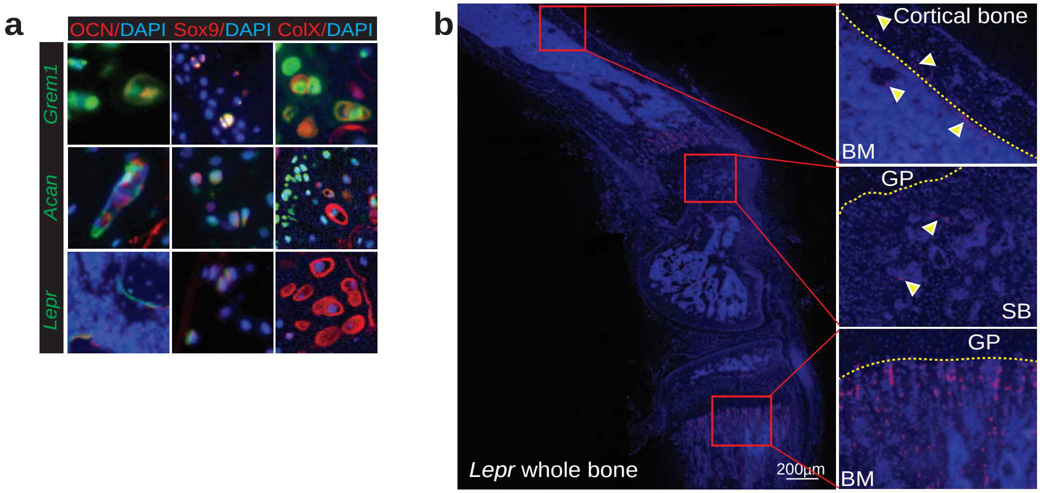


**Figure S2. Related to Figure 2**
**(A)** High magnification images of cells highlighted with yellow arrow in **Figure 2c.**
**(B)** Representative image of whole *LepR-TdT* bone at 8 weeks of age showing lineage tracing in cortical bone, subchondral bone (SB) and bone marrow (BM) just after the growth plate (GP) as indicated by arrows, but not the AC.

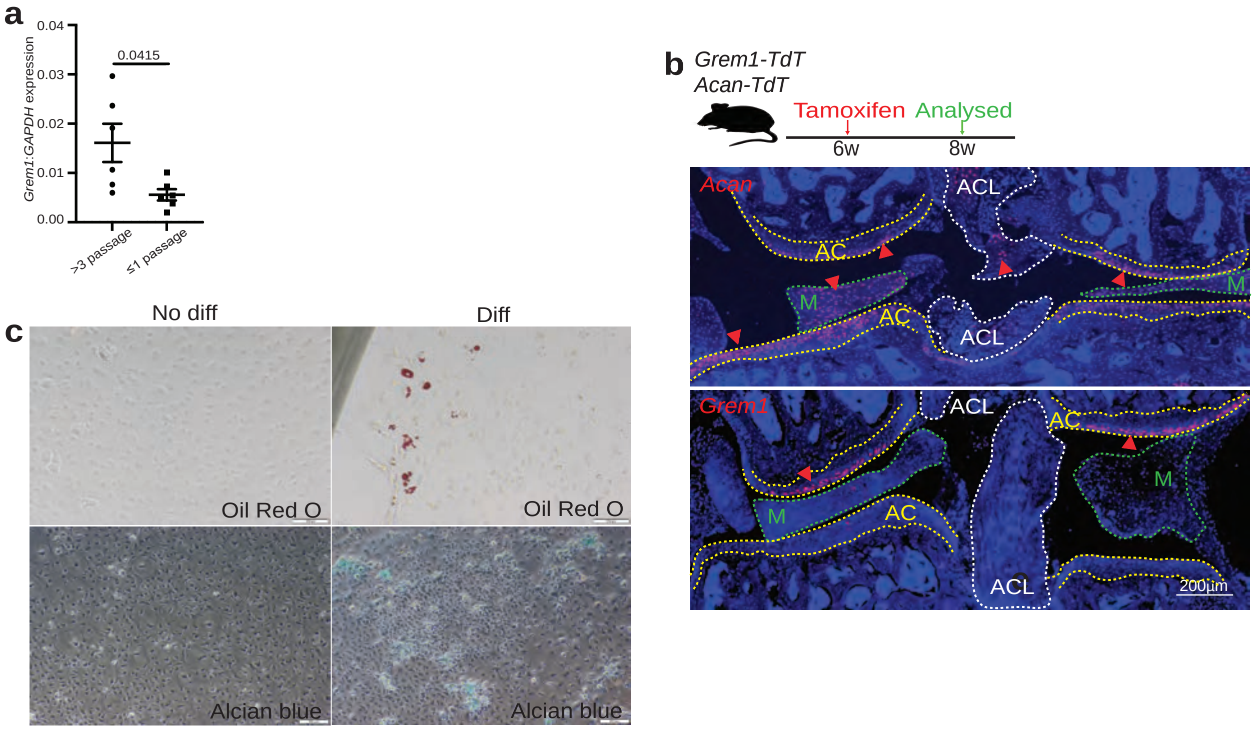


**Figure S3. Related to Figure 3**

**(A)** Real time PCR showed a significantly higher expression of Grem1 in Acan clones that were capable of expansion (●) compared to those that were not (■). Each data point represents an individual clone isolated. Statistical analyses using Welch's t test.

**(B)** Experimental Schema (top). Coronal sections of representative images of *Acan-TdT* (middle) and *Grem1-TdT* (bottom) joint depicting population of lineage traced cells.

**(C)** Osteogenesis and chondrogenesis assays conducted using traditional MSC isolated from whole bone of C57Bl6 animals as positive experimental controls to show successful differentiation into adipogenic and chondrogenic cells using the same media.

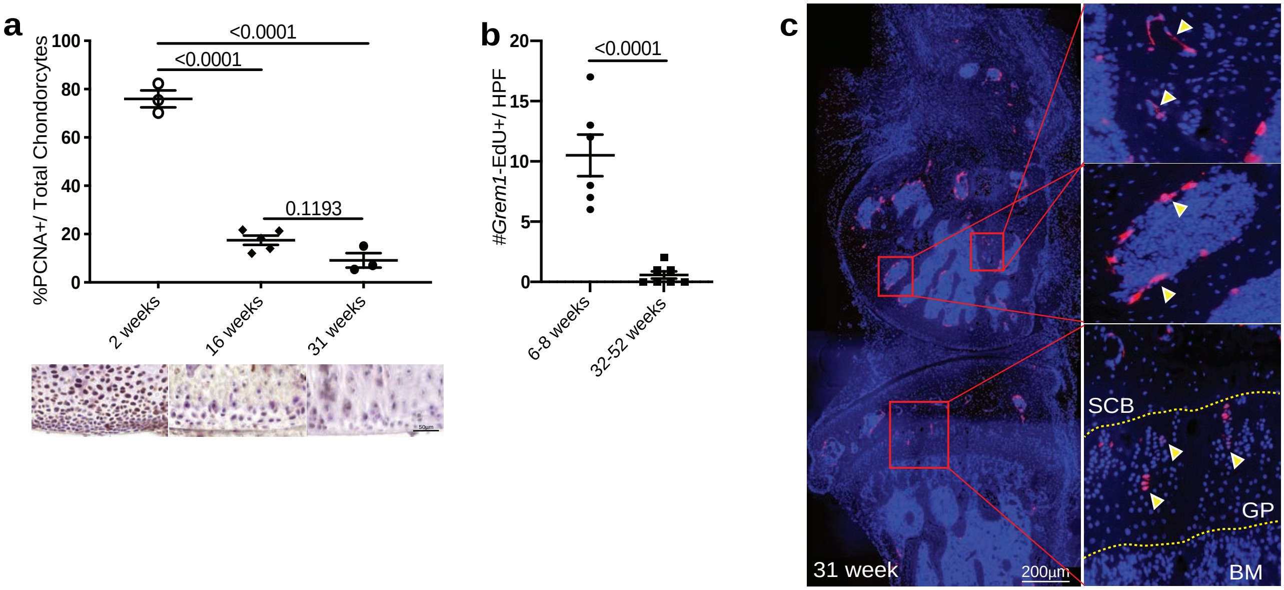


**Figure S4. Related to Figure 4**

**(A)** Quantification of the percentage of PCNA positive cells in the AC at different postnatal stages showed a significant decrease in chondrocyte proliferation from postnatal development to adulthood. Representative images of PCNA staining (bottom). Each data point represents an individual animal analysed, n=3-5 animals per time point. One-way Anova Tukey's test.

**(B)** Quantification of the number of *Grem1*-EdU positive cells of mice in different stages of adulthood showing a significant decrease in proliferation with age. n=6-7 animals per group. Unpaired t test.

**(C)** Representative image of a 31-week-old *Grem1-TdT* bone showing *Grem1*-lineage cells giving rise to osteoblasts within the subchondral bone space (SCB), few chondrocytes within the GP and almost none observed in the AC.

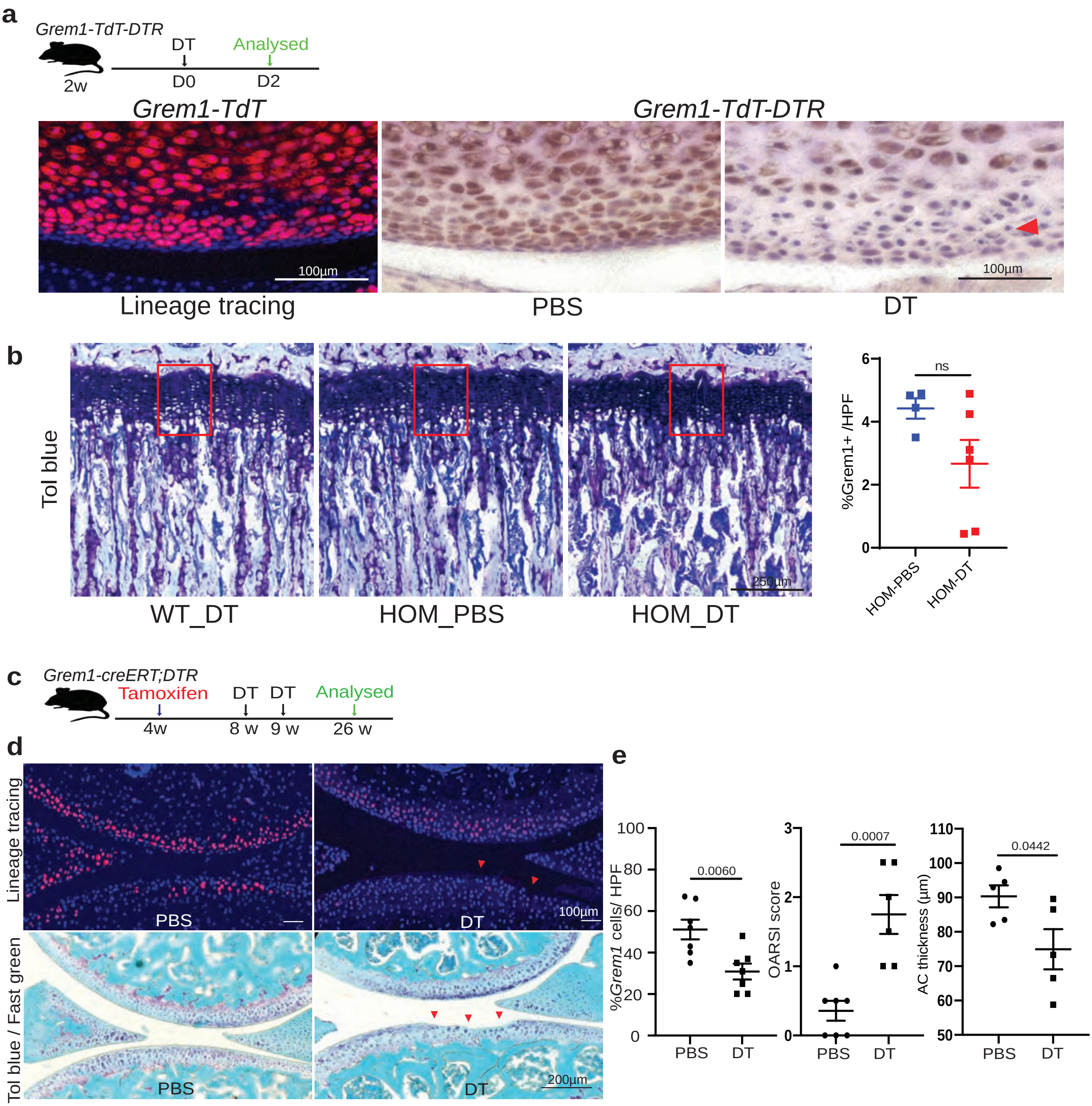

a
Grem1-TdT-DTR
DT
Analysed
2w
D0
D2
Grem1-TdT
Grem1-TdT-DTR
100µm
100µm
Lineage tracing
PBS
DT
b
Tol blue
250µm
WT_DT
HOM_PBS
HOM_DT
%Grem1+ /HPF
ns
6
4
2
0
HOM-PBS
HOM-DT
c
Grem1-creERT;DTR
Tamoxifen
DT
DT
Analysed
4w
8 w
9 w
26 w
d
Lineage tracing
PBS
DT
100µm
Tol blue / Fast green
PBS
DT
200µm
e
%Grem1 cells/ HPF
100
80
60
40
20
0
0.0060
PBS
DT
OARSI score
3
2
1
0
0.0007
PBS
DT
AC thickness (µm)
110
100
90
80
70
60
50
0.0442
PBS
DT

f

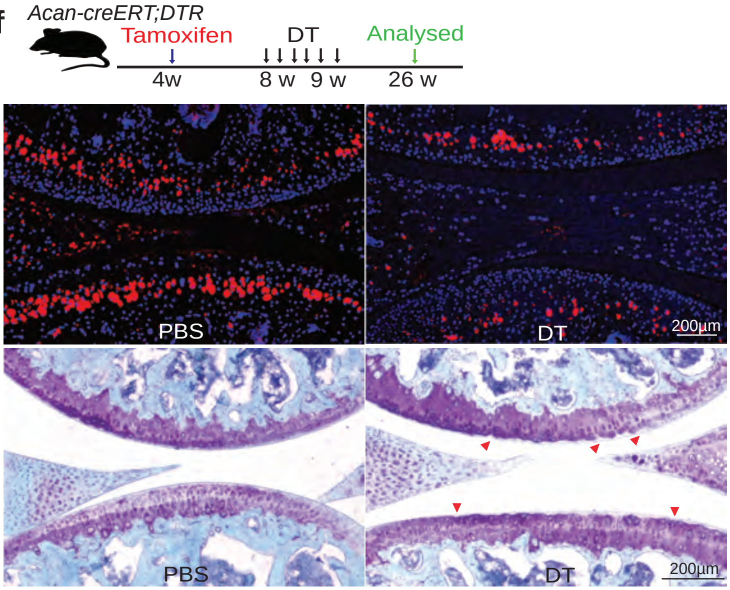

Acan-creERT;DTR
Tamoxifen
DT
Analysed
4w
8 w
9 w
26 w
ACAN
PBS
DT
200µm
Tol blue / Fast green
PBS
DT
200µm


g

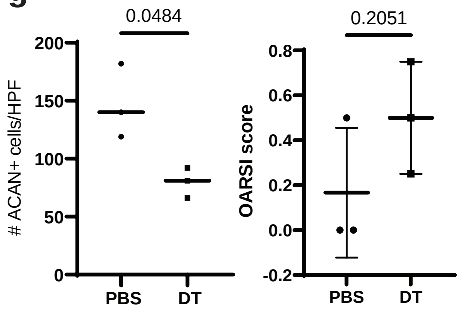

0.0484
### ACAN+ cells/HPF
200
150
100
50
0
PBS
DT
0.2051
OARSI score
0.8
0.6
0.4
0.2
0.0
-0.2
PBS
DT


h

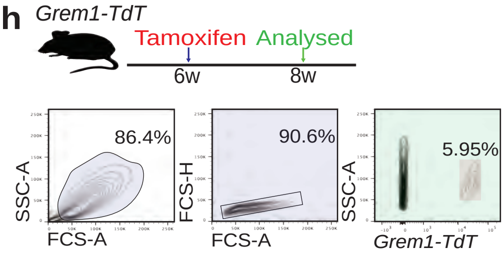

Grem1-TdT
Tamoxifen
Analysed
6w
8w
86.4%
SSC-A
FCS-A
90.6%
FCS-H
FCS-A
5.95%
SSC-A
Grem1-TdT


i

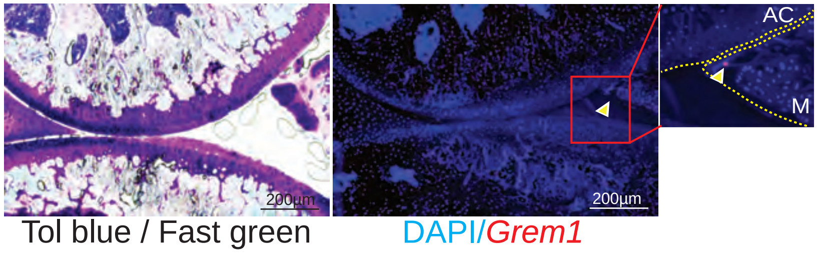

200µm
200µm
AC
M
Tol blue / Fast green
DAPI/Grem1

**Figure S5. Related to Figure 5**

**(A)** Experimental schema (top). Representative images of AC knee joint from 2-week-old lineage traced *Grem1-TdT* mouse (bottom left), in comparison to our new transgenic mouse line *Grem1-Td-DTR* treated with PBS and stained with anti-RFP to visualise *Grem1*-lineage cells (bottom middle) or *Grem1-Td-DTR* mice treated with 250ng of DT also stained with anti-RFP (bottom right). In both *Grem1-TdT* and *Grem1-Td-DTR* mice the *Grem1*-marked cells are distributed identically in the developing joint and targeted ablation of these cells with DT treatment in *Grem1-Td-DTR* mice is indicated with red arrow.

**(B)** Representative images of the growth plate in early adult mice treated with PBS or DT (mice treated as in **Figure 5**) stained with tol blue, showed no significant growth plate changes in sex and age-matched samples. Images from *Grem1-Td-DTR* wildtype (WT) and homozygous (HOM) mice treated with PBS or DT. Quantification of the total *Grem1*-lineage cells in the GP as a percentage of total GP chondrocytes showed no significant depletion with this DT regimen. Welch's t test. n=4-6 mice per group, data point represents an individual animal analysed.

**(C)** Experimental schema.

**(D)** Representative images of AC knee joint from *Grem1-creRET;DTR* mice treated with PBS or DT (top) showing loss of *Grem1* AC cells with DT and stained with Toluidine blue and Fast green with red arrows indicating lesions indicative of OA pathology (bottom). n=7 mice per group.

**(E)** Quantification of the total number of *Grem1* AC cells per HPF showing a significant loss of *Grem1* cells in the AC in DT treated mice (left). Unblinded histopathological assessment using OARSI grading showed a significantly increased OA severity when *Grem1* cells are ablated in the joint compared to PBS control (middle). Quantification of AC thickness in *Grem1-creERT;DTR* mice treated with DT (●) or contralateral joint with PBS (■) as control (right). n=6-7 mice per group, data point represents an individual animal analysed. Unpaired t test.

**(F)** Experimental schema (top). Representative images of AC knee joint from *Acan-creERT;DTR* mice treated with PBS or DT (top) showing loss of ACAN-expressing AC cells with DT, identified using ACAN immunostaining, and stained with Toluidine blue and Fast green with red arrows indicating lesions indicative of OA pathology (bottom). n=3 mice per group.

**(G)** Quantification of the total number of ACAN-expressing AC cells per HPF showing a significant loss of ACAN+ cells in the AC in DT treated mice (left). Unblinded histopathological assessment using OARSI grading showed no significant increase in OA severity when ACAN-expressing cells are ablated in the joint compared to PBS control (right). n=3 mice per group, data point represents an individual animal analysed. Unpaired t test.

**(H)** Experimental schema (top**).** FACS plot showing *Grem1*-lineage cells sorting strategy (bottom).

**(I)** Representative images of DMM knee joints implanted with *Grem1*-lineage sorted cells via intra-articular injection. Serial sections of joint stained with Tol blue and fast green (left) and immunofluorescence image (right) showing *Grem1*-lineage cells homing to the meniscus (yellow arrow).

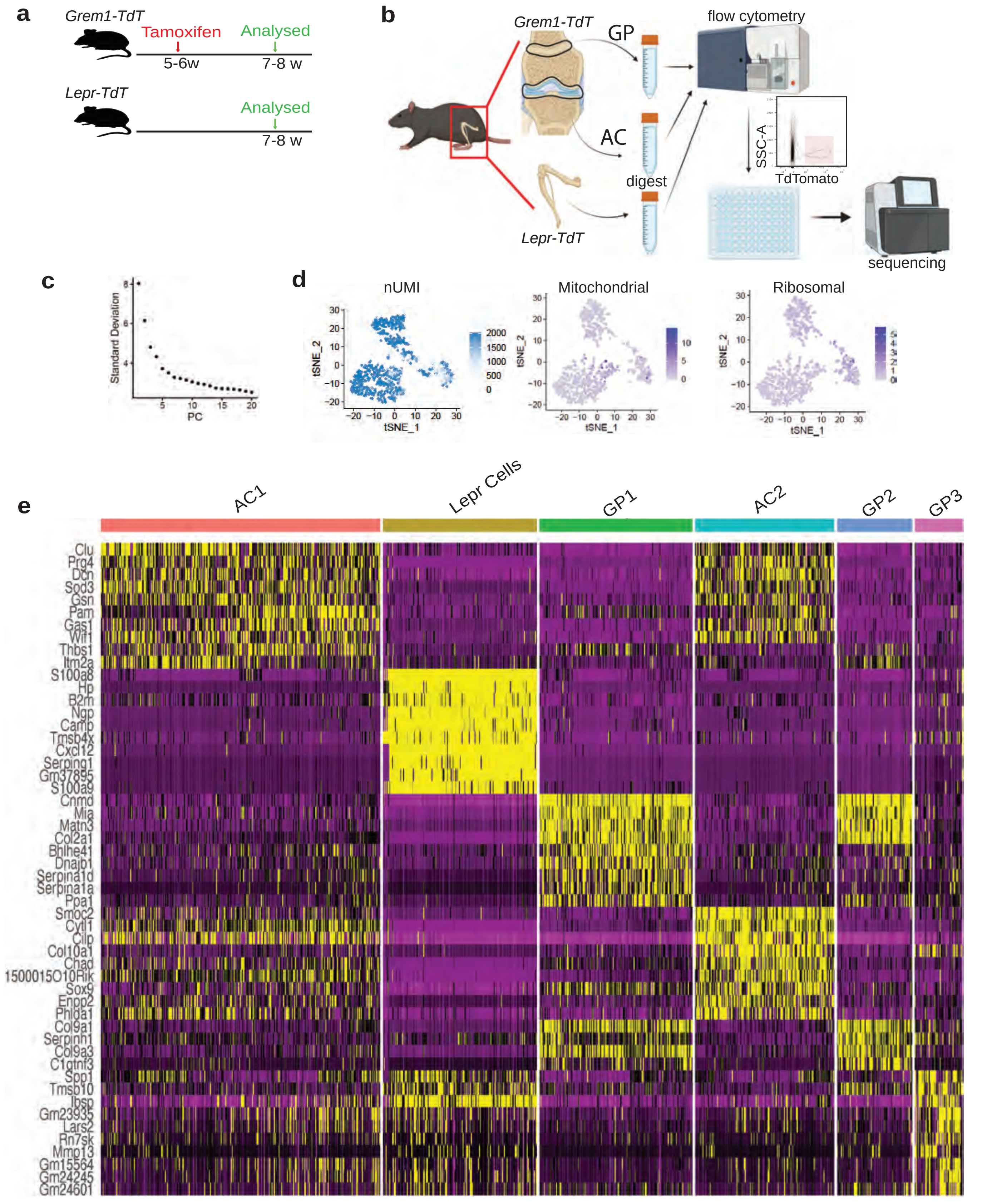
a
Grem1-TdT
Tamoxifen
Analysed
5-6w
7-8 w
Lepr-TdT
Analysed
7-8 w
b
Grem1-TdT
GP
AC
digest
Lepr-TdT
flow cytometry
SSC-A
TdTomato
sequencing
c
Standard Deviation
PC
d
nUMI
Mitochondrial
Ribosomal
tSNE_1
tSNE_2
e
AC1
Lepr Cells
GP1
AC2
GP2
GP3
Clu
Prg4
Dcn
Sod3
Gsn
Pam
Gas1
Wif1
Thbs1
Itm2a
S100a8
Hp
B2m
Ngp
Camp
Tmsb4x
Cxcl12
Serping1
Gm37895
S100a9
Cnmd
Mia
Matn3
Col2a1
Bhlhe41
Dnajb1
Serpina1d
Serpina1a
Pcal1
Smoc2
Cytl1
Clip
Col10a1
Chad
1500015O10Rik
Sox9
Enpp2
Phlda1
Col9a1
Serpinh1
Col9a3
C1qtnf3
Spp1
Tmsb10
Ibsp
Gm23935
Lars2
Rn7sk
Mmp13
Gm15564
Gm24245
Gm24601

f

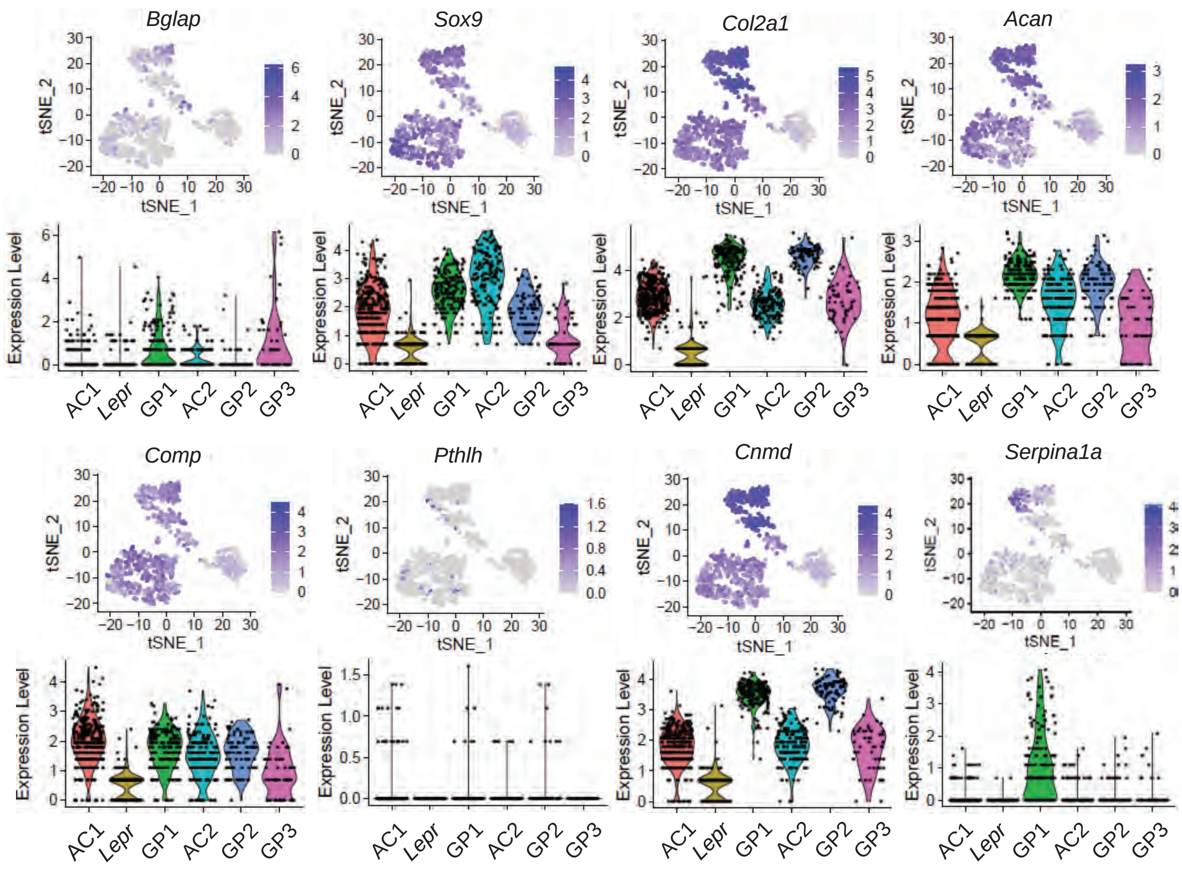
Bglap
Sox9
Col2a1
Acan
Comp
Pthlh
Cnmd
Serpina1a
tSNE_1
tSNE_2
Expression Level
AC1
Lepr
GP1
AC2
GP2
GP3

g

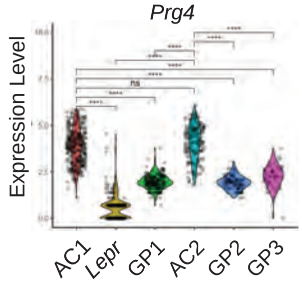
Prg4
Expression Level
ns
AC1
Lepr
GP1
AC2
GP2
GP3

h

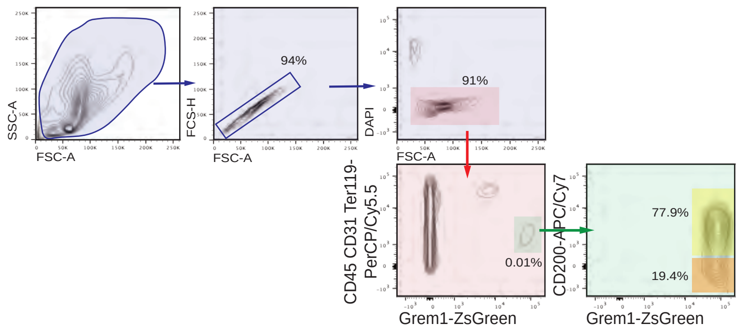

SSC-A
FSC-A
94%
FCS-H
FSC-A
91%
DAPI
FSC-A
CD45 CD31 Ter119-
PerCP/Cy5.5
0.01%
Grem1-ZsGreen
CD200-APC/Cy7
77.9%
19.4%
Grem1-ZsGreen


i

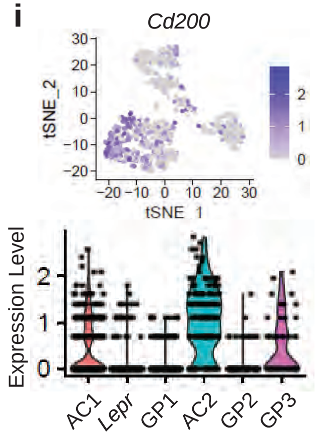

Cd200
tSNE_2
tSNE_1
Expression Level
AC1
Lepr
GP1
AC2
GP2
GP3


i (cont)

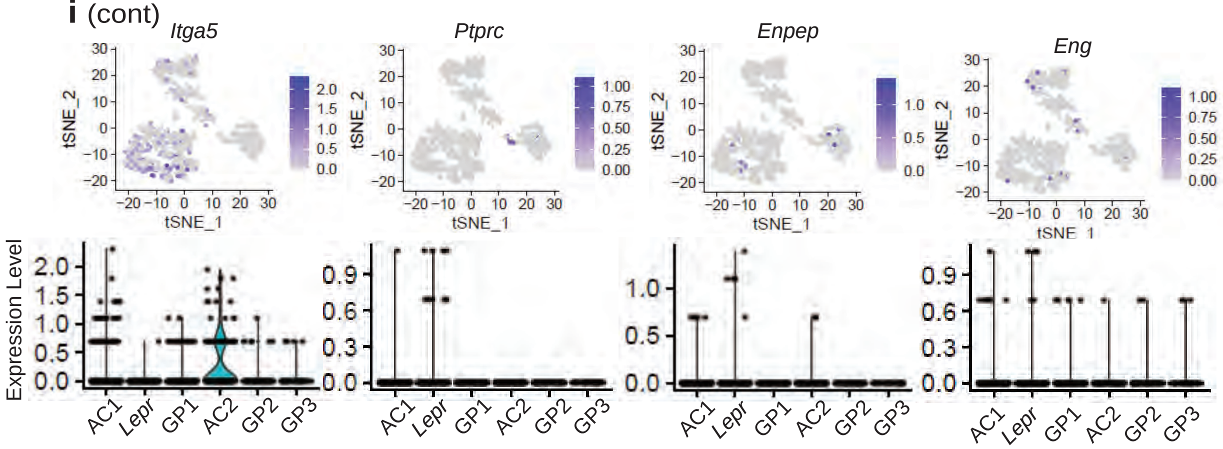

Itga5
Ptprc
Enpep
Eng
tSNE_2
tSNE_1
Expression Level
AC1
Lepr
GP1
AC2
GP2
GP3


j

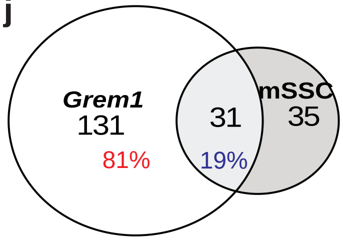

Grem1
131
81%
31
19%
mSSC
35

**Figure S6. Related to Figure 6**
**(A)** Experimental schema showing tamoxifen dosing and analysis schedule.
**(B)** Subsequent dissection of AC, GP or whole bone tissue from *Grem1-TdT* and *LepR-TdT* animals, separately dissociated by digestion, then for each tissue source separately subjected to FACS isolation of TdT positive stem/precursor cell populations, before scRNA sequencing.
**(C)** Elbow plot used to determine appropriate number of principal components (PC) used in scRNAseq data clustering to capture the majority of variation in the data**.**
**(D)** scRNAseq quality control plots showing read depth is appropriate with the majority of unique molecular identifiers (UMI) >1000 (left), low mitochondrial (centre) and ribosomal RNA (right) content.
**(E)** Heat map depicting unsupervised clustering of top 10 differentially expressed transcripts between the different scRNAseq clusters as shown in Figure 6 but here displayed with gene names.
**(F)** Expression of classical osteoblast (osteocalcin, *Bglap*), chondrocyte (*Sox9, Col2a1, Acan, Comp, Cnmd*) and growth plate markers (*Pthlh*) in the scRNAseq dataset.
**(G)** Expression of AC marker, *Prg4* (encoding Lubricin), is significantly increased in the two AC clusters in comparison to the GP and *Lepr* cell populations, **** $p<0.0001$ Tukey's test.
**(H)** Immunophenotyping of *Grem1*-lineage cells from total bone preps of 8-week-old mice using flow cytometry showed that 77.9% express the key mSSC marker CD200. n=3 animals.
**(I)** tSNE and violin plots depict expression of transcripts encoding additional immunophenotypic markers of AlphaV$^{+}$CD200$^{+}$CD45$^{-}$6C3$^{-}$CD105$^{-}$Ter-119$^{-}$Tie2$^{-}$Thy$^{-}$ mSSC cells in the scRNAseq data from *Grem1*- and *Lepr*-lineage cells. *Itga5* encodes AlphaV, *Ptprc* encodes CD45, *Enpep* encodes 6C3, *Eng* encodes CD105, while transcripts for *Ly76* (encoding Ter119), *Tek* (encoding Tie2) and *Thy1* (encoding Thy) were not detected.
**(J)** Of the *Grem1*-lineage AC population that express *Grem1* in the scRNAseq data, only 19% also expressed mSSC immunophenotype genes, indicating most *Grem1*-expressing AC cells are likely distinct from mSSCs.

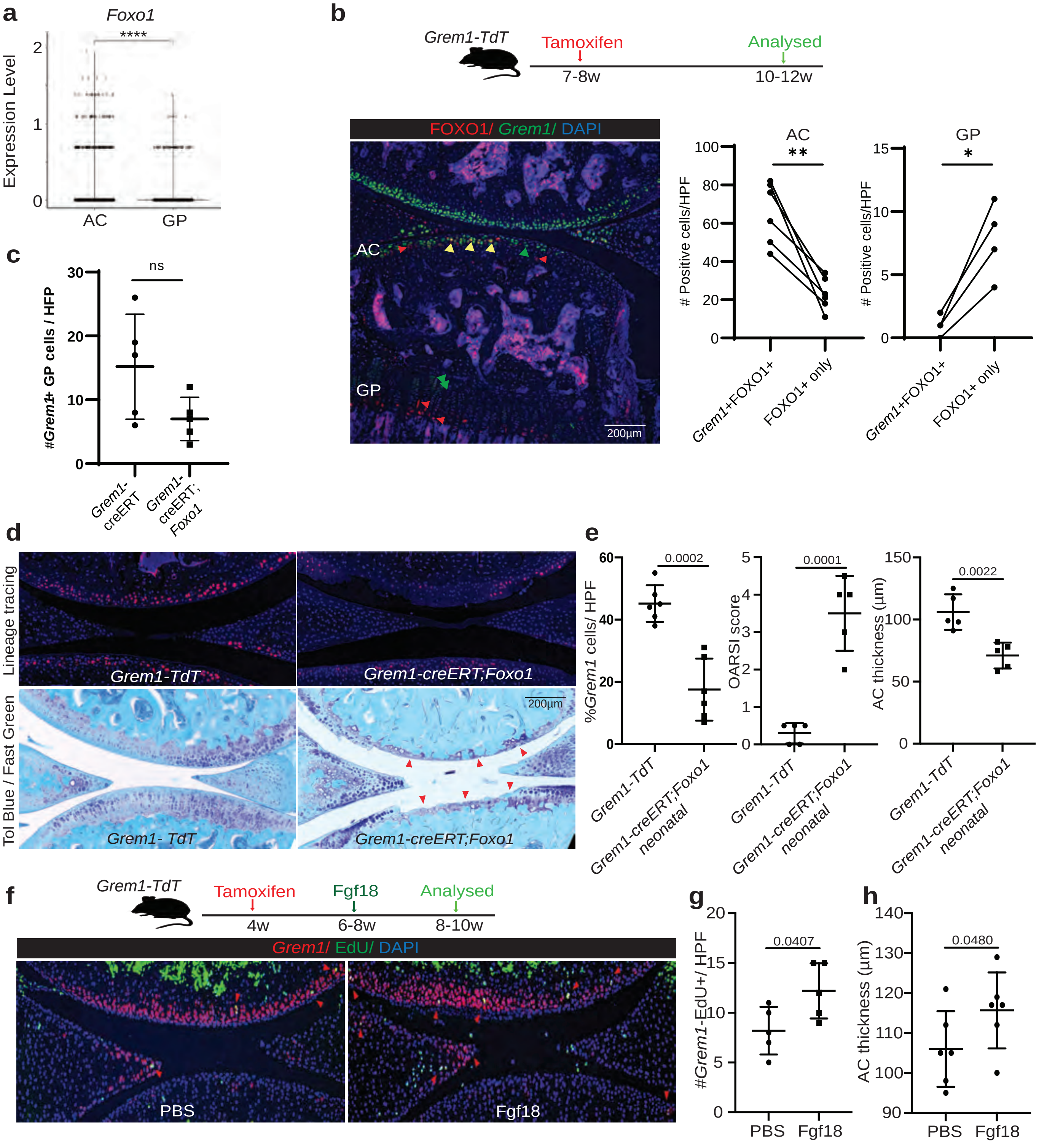

a
Foxo1
****
Expression Level
0
1
2
AC
GP
b
Grem1-TdT
Tamoxifen
Analysed
7-8w
10-12w
FOXO1/ Grem1/ DAPI
AC
GP
200µm
AC
**
GP
*
### Positive cells/HPF
Grem1+FOXO1+
FOXO1+ only
c
ns
#Grem1+ GP cells / HFP
Grem1-creERT
Grem1-creERT;Foxo1
d
Lineage tracing
Tol Blue / Fast Green
Grem1-TdT
Grem1-creERT;Foxo1
Grem1- TdT
200µm
e
0.0002
0.0001
0.0022
%Grem1 cells/ HPF
OARSI score
AC thickness (µm)
Grem1-TdT
Grem1-creERT;Foxo1 neonatal
f
Grem1-TdT
Tamoxifen
Fgf18
Analysed
4w
6-8w
8-10w
Grem1/ EdU/ DAPI
PBS
Fgf18
g
0.0407
#Grem1-EdU+/ HPF
h
0.0480
AC thickness (µm)
PBS
Fgf18

**Figure S7. Related to Figure 7**

**(A)** Expression of *Foxo1* in pooled articular cartilage (AC) or growth plate (GP) *Grem1*-lineage populations in scRNAseq dataset (described in **Figure 6, Extended data Fig. 6**). Tukey's test, **** p<0.0001.

**(B)** Experimental schema and representative image of FOXO1 expression (red) in *Grem1*-lineage (green) cells in the AC but not GP. Red arrows indicate FOXO1+ only cells, green arrows *Grem1*-lineage only, yellow arrows both FOXO1+ and *Grem1*-lineage cells. Quantification of fluorescently labelled average cell number in the AC and GP to the right, n=6 mice per group, 2-4 HPF/animal; t-test, *p<0.05, **p<0.01.

**(C)** Quantification of the number of *Grem1*-lineage cells in the GP of *Grem1-TdT-Foxo1* deletion mice compared to *Grem1-TdT* mice showed no significant (ns) difference, schema for experiment depicted in **Fig 7a.** n=5 mice per group, 2-3 HPF/animal; t-test.

**(D)** Representative images of neonatally induced deletion of *Foxo1* in *Grem1-TdT* cells showing significant loss of AC (arrows).

**(E)** Quantification of the percentage of *Grem1*-lineage articular chondrogenic progenitor (CP) cells per HFP showed a significant decrease in *Grem1*-lineage CP cells in neonatal induced *Grem1-TdT-Foxo1* mice (■) compared to *Grem1-TdT* mice (●) (left). Unblinded histopathological scoring of OA pathology also showed a significant increase in OARSI score in *Grem1-TdT-Foxo1* deletion mice (■) compared to *Grem1-TdT* mice (●) (middle), and AC thickness indicating the important role of *Grem1*-lineage CP cells in early AC development through *Foxo1* signalling (right). Each data point represents an individual animal analysed, n=5 mice per group.

**(F)** Experimental schema (top). Representative IF image of EdU staining showing an increase in proliferating *Grem1*-lineage articular CP cells indicated by red arrows with FGF18 treatment (bottom).

**(G)** Quantification of the number of EdU positive *Grem1*-lineage articular CP cells showed that FGF18 treatment (■) significantly increased *Grem1*-lineage articular CP cell proliferation compared to PBS control (●).

**(H)** This increase in proliferation likely resulted in the increase in AC thickness in *Grem1*-TdT mice treated with FGF18 (■) compared to contralateral knee treated with PBS as control (●). Statistical analyses used Unpaired t test.

**a**

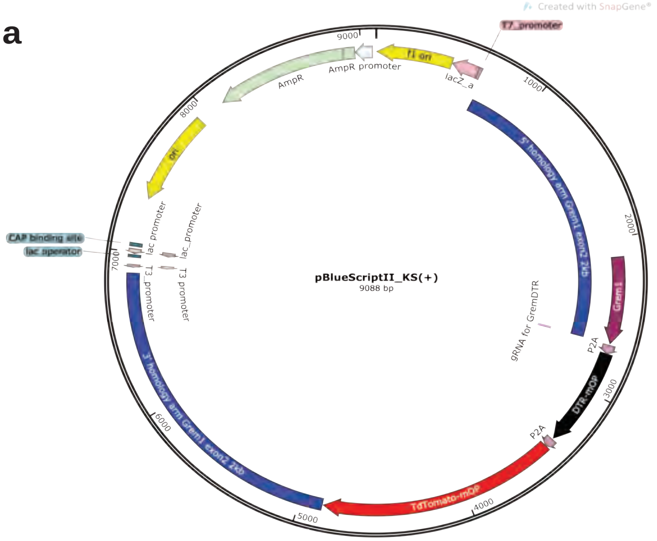


**Figure S8. Diagram of plasmid construct for generation of Grem1-TdT-DTR knock-in mouse strain.**

| Cluster | Top 10 Cluster defining transcripts |
|---|---|
| AC1 | *Clu,Prg4,Sod3, Gsn, Pam, Itm2a, Cytl1,Cilp,Chad, Phlda1.* |
| LepR | *S100a8, Hp,B2m, Ngp, Camp,Tmsb4x, Cxcl12,Serping1,Gm37895,S100a9.* |
| GP1 | *Cnmd,Mia,Matn3,Col2a1,Bhlhe41,Dnajb1,Serpina1d, Serpina1a,Ppa, Col9a1.* |
| AC2 | *Smoc2,Cytl1,Cilp,Col10a1,Chad,1500015O10Rik,Sox9,Enpp2,Phlda1,Prg4.* |
| GP2 | *Col9a1,Serpinh1,Col9a3,Cnmd,C1qtnf3,Col2a1,Matn3,Mia,Eno1b,Eno1.* |
| GP3 | *Spp1,Gm23935,Lars2,Rn7sk,Gm23037,Bglap3,Bglap2,Bglap,Col1a1,Col1a2.* |

**Table S1. Top 10 transcripts that define each cluster in the scRNAseq dataset.**
