## Supplementary material for "Loss of *Grem1*-articular cartilage progenitor cells causes osteoarthritis": Methods file

**STAR Methods**

**RESOURCE AVALIBILITY**

**Lead Contact**

Further information and requests for resources and reagents should be directed to and will be fulfilled by the lead contacts, Siddhartha Mukherjee and Susan Woods.

**Materials Availability**

Mouse line generated through this study has not been made available to the public. Any information required is available from the lead contacts upon request.

**Data and Code Availability**

**Data**

Single-cell RNA-seq data have been deposited at GEO and are publicly available as of the date of publication with the accession number GSE193175. Data reported in this paper will be shared by the lead contacts upon request.

**Code**

This paper does not report original code.

Any additional information required to reanalyse the data reported in this paper is available from the lead contacts upon request.

**EXPERIMENTAL MODEL AND SUBJECT DETAILS**

**Mice**

*LepR-cre(DeFalco et al., 2001)*, *Acan-creER^T2(Henry et al., 2009)^*, *Grem1-creER^T(Worthley et al., 2015)^*, *R26-LSL-TdTomato(Madisen et al., 2010)*, *R26-LSL-ZsGreen(Madisen et al., 2010)* and *R26-LSL-iDTR* were imported from Jackson Laboratory and bred within the South Australia Health and Medical Research Institute (SAHMRI) Bioresources facility and Institute of Comparative Medicine at Columbia University. *Grem1-TdTomato-iDTR* mice (*Grem1-creERT;DTR)* were generated by mating homozygous *Grem1-TdT* mice to homozygous *Rosa-iDTR*. *Acan-iDTR* mice (*Acan-creERT;DTR)* were generated by mating heterozygous *Acan-creERT* mice to homozygous or heterozygous *Rosa-iDTR*. All animal experiments complied with all relevant ethical regulations and were approved by the Animal Ethics Committee at the SAHMRI under ethics number SAM189 and SAM385.19 and at Columbia University.

Mice were maintained in the South Australia Health and Medical Research Institute (SAHMRI) Bioresources facility and Institute of Comparative Medicine at Columbia University, in accordance with JAX USA animal husbandry protocols. Animals were given food and water ad libitum and housed in temperature-, moisture-, and light-controlled (12h light/dark cycle) individually ventilated cage systems. 10 – 13 weeks old *Grem1-TdT,* *Acan-TdT, LepR-TdT* and *C57BL/6* male mice were used to examine lineage tracing in health and DMM surgery model. Postnatal Day 4 – 6 (neonatal) and 6 weeks old (adult) *Grem1-TdT, Acan-TdT* and *LepR-TdT* mice were used to examine lineage tracing in early development and adulthood. 8 weeks old Grem1-TdT mice were given tamoxifen at 6 weeks of age to examine in vitro stem cell properties. 26 week old *Grem1-creERT;FOXO1* mice were given tamoxifen at postnatal Day 4 – 6 (neonatal) and 6 weeks of age (adult) to examine the AC impact of *FOXO1* deletion in *Grem1* cells. 4 – 6 week old *Grem1-TdT* mice were used to examine the treatment of Fgf18 in AC with or without CiOA. 4 – 7 week old *Grem1-Td-DTR*, *Grem1-creERT;DTR* and *Acan-creERT;DTR* mice were used to examine the impact of the ablation of *Grem1*-lineage and *Acan*-lineage cells on AC.

**Destabilisation of the medial meniscus surgery (DMM)**

DMM surgery was conducted using an established protocol(Glasson et al., 2007) approved by the SAHMRI Bioresources Animal Ethics Committee. Briefly, surgery was performed on male mice ­­between the ages of 10 – 13 weeks that had previously been administered tamoxifen. Animals underwent general anaesthesia via isoflurane inhalation. The right hind limb of the animal was shaved with a razor blade to provide a clean surface and left hind limb was used as a paired normal control. Skin incision was made on the medial side of the patella, using sterile scissors, to expose the patella. Once the patella was exposed, a second incision was made on the medial side and the incision site was extended so that the patella could be luxated. Using sterile gauze, the fat pad located on the knee was pushed aside to expose the medial meniscotibial ligament, the ligament was then transacted using a twisting motion. Bleeding was controlled throughout surgery, and the cartilage was kept moist with sterile saline. The patella was then repositioned back, and the incision sites and patella were sutured.

**Collagenase VII induction of OA (CiOA)**

The collagenase-induced OA model involves intra-articular injection of Type VII Collagenase as described(Botter et al., 2008). Briefly, 8-week-old mice previously administered with tamoxifen were injected intra-articularly with 10µL of 2U ColVII (#C0773, Sigma-Aldrich) dissolved in PBS under a dissection microscope.

**Generation of knock-in *Grem1*-DTR-TdT mice by CRISPR/Cas9 mediated genome engineering**

To generate a DNA construct for Diphtheria Toxin Receptor (DTR) / tdTomato (tdTom) expression under the control of the endogenous *Grem1* promoter in mice, we first used gene synthesis to insert the coding sequence for DTR immediately downstream of a 2A peptide from porcine teschovirus-1 polyprotein (P2A), followed by a second P2A sequence and the coding sequence for tdTom, all within two exon2 2kb *Grem1* homology arms (**Figure S8**). The construct was Sanger sequence verified (Geneuniversal) and used for subsequent gene targeting in mice. C57BL6/J female mice were superovulated with 5 IU Pregnant Mare Serum Gonadotrophin (PMSG; Folligon; Intervet) and 47.5 hours later with 5 IU human chorionic gonadotrophin (hCG; Chorulon; Intervet) before being mated to C57BL6/J males. Presumptive zygotes were collected from oviducts 19 hours post-hCG injection in FHM media (Merk Millipore) and maintained in KSOMAA media (Merk Millipore) under oil at 37 °C in 5% CO_2_ 5% O_2_ with a Nitrogen balance. Fertilized zygotes were identified in FHM media under oil by the presence of two pronuclei before microinjection of 25 ng/ul SpCas9 protein (PNA Bio), and 10 ng/ul SpCas9 single guide RNA (sgRNA) (CGACTTGGATTAAGTCAAAG) with the donor plasmid at 10ng/ul, in a buffered solution. Genomic DNA was isolated from ear notches using a High Pure PCR Template Preparation Kit (Roche) and Knock-in (KI) founder screening was performed using a forward primer (GCCTGGGATATTCTGAGCCC) and a reverse primer (GTCCCGCGCATTTTGACTTT). KI allele in the founder mouse was confirmed by PCR and Sanger sequencing across the entire locus from outside of the donor homology arms before mating with C57BL6/J to create second generation line.

**METHOD DETAILS**

**Tamoxifen administration**

Neonatal *Grem1-TdT* and *Acan-TdT* mice were subcutaneously administered 2mg tamoxifen (#T5648, Sigma) dissolved in peanut oil at Day 4 – 6 of age for lineage tracing studies. *Grem1-TdT* and *Acan-TdT* adult lineage tracing mice were administered 4 doses of 6mg tamoxifen dissolved in peanut oil via oral gavage at ages specified in figures. *Grem1-TdT* and *Acan-TdT* DMM OA mice were administered 4 doses of 6mg tamoxifen dissolved in peanut oil via oral gavage at 8 – 10 weeks of age. *Grem1-TdT* ColVII OA, *Grem1-creERT;DTR* and *Acan-creERT;DTR* mice were administered 5 separate doses of 4mg tamoxifen via daily oral gavage over 10-14 days from 4-6 weeks of age. EdU labelling of *Grem1-TdT* mice was performed by administering 5 separate doses of 4mg tamoxifen via daily oral gavage over 9 days and commenced on EdU in drinking water 2 weeks prior to collection.­­ *Grem1-TdT* mice for scRNA experiments were administered 500mg/kg tamoxifen chow for 2 weeks from 5 – 6 weeks of age.

**Diphtheria toxin (DT) administration**

*Grem1* cells within the AC are abundant at 2 weeks of age, therefore, we selected mice at this age for initial validation of *Grem1* cell ablation within the AC. Diphtheria toxin (#BML-G135, Enzo) at a concentration of 250ng in 10µL of PBS was injected intraperitoneally into 2-week-old *Grem1-Td-DTR* and mice sacrificed 2 days later. Anti-RFP immunohistological staining was performed to validate the distribution and ablation of *Grem1* cells in the *Grem1-Td-DTR* AC compared to aged matched *Grem1-TdT* mice (**Figure S5A**). For OA pathology, mice 5 – 7 weeks of age were administered two doses of 250ng DT intra-articularly in 10µL of PBS and limbs collected 3 days later for analysis and blinded scoring of OA pathology. *Grem1-Td-DTR* mice administered with DT developed a fatal small intestinal phenotype at day 4 for homozygous mice and at day 7 for heterozygous mice after DT injection. A similar phenotype has been previously reported(McCarthy et al., 2020). Ablation of *Grem1*-lineage cells in *Grem1-creERT;DTR* mice and *Acan*-lineage cells in *Acan-creERT;DTR* mice was performed by administering 100-125ng of DT (#D0564, Sigma) intra-articularly 3 times weekly for 2 weeks into 8 week old mice that were given tamoxifen prior at 4-6 weeks of age. Mice were then sacrificed at 26 weeks of age for analysis and unblinded OARSI scoring.

***Grem1* Chondroprogenitor Cell isolation**

Both hind limb bones (femur and tibia) were collected from 8-week-old mice that had been subjected to tamoxifen and all muscles and ligaments removed. To limit contamination from non-articular cell populations present in whole bone preparations, the knee joints were dislocated at the epiphyses from both femur and tibia to collect the articular region only, gently disrupted using a mortar and pestle, and digested in 4mg/ml collagenase IV (#17104019, Gibco) and 3mg/ml Dispase (#17105041, Gibco) in α-MEM (#M5650, Sigma). Collected cells were then sorted for lineage-traced articular *TdTomato* positive cells. Cells were sorted based on forward and side scatter, single cells as well as positive for *TdTomato* fluorescence. Live and dead (DAPI) staining was omitted to allow for later expansion of clones in DAPI free conditions. Sorted cells were plated in complete media ( α-MEM supplemented with 20% defined bovine serum, 100mM L-ascorbate-2-phosphate, 1mM sodium pyruvate, 50μg/ml streptomycin, 50U/ml penicillin, 2mM L-glutamine and 10μM Y-27632) at 1000 cells per 10cm dish to allow for isolation of individual clones using clonal cylinders (#Z370789, Sigma). A total of 22 – 24 clones (i.e., isolated adherent clusters of >50 cells at 10-14 days) were harvested and allowed to expand before performing clonogenicity and multilineage differentiation assays. Both clones that expanded and did not expand were collected for qPCR analysis.

**Clonogenicity and multilineage differentiation**

Colony-forming unit – fibroblasts (CFU-F) assay was performed by seeding cells at clonal density (1000 cells per 10cm dish) and subsequently cultured in complete media for 14 days. The cultures were stained with 0.1%w/v toluidine blue in 2.4% formalin solution and the total number of colonies counted, where an individual colony was defined as ≥50 cells. The number of clones was reported as (CFU-F)/1,000 cells plated. Induction of osteogenic, chondrogenic and adipogenic differentiation was conducted by maintenance of cell cultures in differentiation media (StemPro) as per manufacturer’s instructions. Positive and negative assessment of differentiation potential was performed by staining cells with 2% w/v Alizarin red in water pH 4 (osteogenesis), 0.1% w/v Alcian blue in 0.1N HCl (chondrogenesis) and 0.5% w/v Oil Red O in isopropanol diluted further in a 6:4 ratio in water (adipogenesis). All *in-vitro* assays reported in this study were conducted on clonal cell populations of passage 5 or lower.

**Histology**

Mice were humanely sacrificed, bones from both hind limbs were collected and all muscles removed before fixing in 4% paraformaldehyde overnight, decalcified in Osteosoft® (#101728, Millipore) for 3 – 10 days and dehydrated in 30% sucrose at 4°C before embedding in OCT compound (Sakura Tissue-Tek). Embedded tissues were stored at -80°C. 10μm frozen sections were collected on cryofilm (type IIC(10), Section-Lab) for staining. 0.04% toluidine blue (#198161, Sigma) in 0.1M sodium acetate pH4.0 or 0.1% Safranin O (#8884, Sigma) in water, and 0.1% fast green (#F7252, Sigma) stain in MilliQ water was used to demonstrate histological features of cartilage and bone. Haematoxylin and eosin (H&E) staining of the AC region was used to identify different organisational zones within the adult AC.

**Immunohistological and fluorescent staining**

Immunohistochemistry and immunofluorescent staining were completed on 10μm frozen sections. Antigen retrieval was performed by placing slides in a steamer submerged in antigen unmasking solution (#H-3300, VectorLab) for 6 min. 0.025% v/v triton X-100 in PBS was used for anti-RFP staining. Immunohistochemistry slides were treated for endogenous peroxidase activity by incubating in 3% H_2_O_2_ for 30 min. Blocking was performed in 2% BSA, 5% normal goat and 5% normal donkey serum. The following antibodies were used: anti-PCNA (#ab18197 Abcam, 1:200), ColX (#ab58632 Abcam 1:200), OCN (#ab93876 Abcam, 1:200), Lubricin/PRG4 (#ab28484 Abcam, 1:250), COL2 (#ab34712 Abcam, 1:250), Sox9 (#AB5535 Millipore, 1:400), ACAN (#AB1031 Merck Millipore, 1:100) and anti-RFP (#600-401-379, Rockland 1:250). After overnight incubation at 4°C, slides were washed with PBST and incubated with species-appropriate secondary antibody (1:200-1:300) at room temperature for 1 h. Finally, slides were counter stained with DAPI before mounted with a cover slip. For immunohistochemistry, after overnight incubation at 4°C, slides were washed with PBST and incubated with anti-rabbit biotin (#BA-1000, VectorLab 1:250) at room temperature for 1 h and then streptavidin-HRP (#SA-5004, VectorLab 1:100) at room temperature for 30 min and developed with DAB chromogen (#K3468, Dako). EdU staining was done using the Click-iT EdU proliferation kit (#C10086, ThermoFisher) as per manufacturer’s instructions.

**Imaging**

Stained sections were scanned using the 3DHistech Panoramic 250 Flash II to generate brightfield images. Fluorescent images were captured either on a Olympus IX53 and Nikon Eclipse Ti inverted microscope or the Leica TCS SP8X/MP confocal microscope. Images of whole bone were achieved by stitching together 20 – 24 images at 10x magnification using ImageJ software.

**RNA isolation and RT-PCR**

Total RNA was isolated from cells using TRIzol (Thermofisher) as per manufacturer’s instructions. Complementary DNA (cDNA) was generated using SuperScript IV reverse transcriptase kit (Invitrogen) according to manufacturer’s protocol. Transcript levels were assessed by QuantStudio 7 (Thermofisher) using IDT probes GAPDH (Mm.PT.39a.1) and Grem1 (Mm.PT.53a.31803129) with the FastStart TaqMan Probe Master (#04673409001, Roche).

**Flow Cytometry for assessment of mSSC markers in *Grem1*-lineage cells**

All limbs were collected from 8-week-old mice that had been subjected to tamoxifen and all muscles and ligaments removed. Cleaned bones (i.e. whole bone preparation) were gently disrupted using a mortar and pestle, minced, and digested in 2.5mg/mL collagenase I (#CLS-1, Worthington). Collected cells were subjected to RBC lysis and blocked with 3% BSA, 2% FCS in PBS. The following antibodies were used: anti-CD45 (#103131, BioLegend), anti-Ter119 (#116227, BioLegend), anti-CD31 (#562861, BD Pharmingen), anti-CD200 (#ab33735, Abcam), APC/Cy7 Streptavidin (#405208, BioLegend) and DAPI (#D9542, Sigma-Aldrich). Cells were sorted on FACSFusion or LSR Fortessa (BD Biosciences) based on forward and side scatter plots, single cells, live cells (DAPI negative), trilineage expression (CD45^-^Ter119^-^CD31^-^) and lineage reporter positive fluorescence.

**Single cell sorting for scRNAseq analysis**

*Grem1-TdT* mice at 5 – 6 weeks of age were subjected to tamoxifen chow for 2 weeks before they were sacrificed. Age paired *LepR-TdT* mice were used for this analysis. Hind limbs were collected, and all muscles and ligaments removed. AC and GP from *Grem1-TdT* mice were separately excised under a dissection microscope and kept as separate tissue samples (AC and GP) through subsequent mincing using a scalpel before being digested in 2.5mg/mL collagenase type II (#CLS-2, Worthington). Cleaned *LepR-TdT* whole bones were gently disrupted using a mortar and pestle, minced, and digested in 2.5mg/mL collagenase type I (#CLS-1, Worthington). The following antibodies were used: anti-CD45 (#103111, BioLegend), anti-Ter119 (#116211, BioLegend), anti-CD31 (#102509, BioLegend), and DAPI (#D9542, Sigma-Aldrich). *LepR* cells were sorted based on forward and side scatter, single cells, live cells (DAPI negative), trilineage negative expression (CD45-Ter119-CD31-) and TdTomato positive fluorescence. *Grem1* cells from the AC or GP were sorted separately based on forward and side scatter, single cells, live cells (DAPI negative), and TdTomato positive fluorescence, to generate separate *Grem1*-lineage populations derived from the AC or GP. Individual cells were collected into single well in a 96-well plate with lysis buffer previously prepared by the Sulzberger Genome Center (Columbia University) and sequenced. Cells were randomly allocated into 96-well plates to avoid batch effects.

**QUANTIFICATION AND STATISTICAL ANALYSIS**

**Quantification of cells and AC thickness**

Lineage-traced (*TdTomato* positive) cells within the AC were counted throughout the femur and represented as a percentage of the total number of chondrocytes (DAPI positive). Only *TdTomato* cells colocalised with DAPI were included to avoid the inclusion of dead cells. AC cells in neonatal tracing were defined by the thin and highly cellular structures consisting of small and closely bound cells. Serial sections stained with H&E were used to guide the zonal separation of the AC. Cells in both the femur and tibia AC in 3 randomly selected images at 40x magnification per sample were used to quantify the number of lineage-traced cells per high power field (HPF) and represented as a percentage of the total number of chondrocytes. AC thickness was measured by averaging 3 randomly selected areas of the AC in the femur and tibia. Final AC thickness was represented as an average of 3 slides per sample. EdU and TUNEL positive *Grem1*-lineage cells were quantified in 3 randomly selected images at 40x magnification per sample and represented as total number of positive cells per HPF. Only EdU positive cells within 1-4 cells from the superficial zone of the articular layer were counted.

**OA pathology scoring**

OA pathology (both femoral and tibial) in *Grem1-creERT;DTR* mice was scored using histological sections in an unblinded fashion using the 0 – 6 OARSI scoring system(Glasson et al., 2010). Final OARSI scores represent the average of 3-6 slides per sample. OA pathology in the *Grem1-TdT-DTR* experiments were scored blinded using additional scoring parameters such as AC damage, proteoglycan loss, chondrocyte hypertrophy, subchondral bone invasion and meniscus pathology. Each of these parameters were scored and reported separately using a 0 – 3 scoring paradigm where 0 is normal, 1 = mild, 2 = moderate and 3 = severe changes. The average scores of 2 – 3 slides per sample, total of 5 parameters, were summed to produce the final score of 0 – 15. The average OA scores of both tibia and femora were included in this study.

**Single Cell Sequencing analysis**

Cells from 2 – 3 individual samples were grouped together, and count tables were analysed in the R environment (version 4.0.2) with the Seurat package (version 3.2.2). All lineage traced cells were combined into one single cell object. When using FindClusters() function, dimensions of reduction were set from 1 to 9 and resolution was set at 0.6 to capture the majority of variation in the data (**Figure S6C)**. This unsupervised clustering identified 6 distinct cell clusters. We undertook quality control assessments to check that sequencing depth (indicated by unique molecular identifiers per cell), sample quality (mitochondrial and ribosomal content per cell) and proliferation markers were not drivers of cluster identity (**Figure S6D).**

To explore single cell clusters that co-express *Grem1*, *Foxo1* and *Fgfr3 a g*ene count greater than 0 was used to define positive expression, and Chi-Square statistics were employed. For cluster analysis, FindAllMarkers() was used to generate marker lists, with parameters of min.pct = 0.25, logfc.threshold = 0.25, and tes.use = “roc”. To investigate the relationship between *Grem1* AC cells with previously published mouse skeletal stem cells(Chan et al., 2015) (mSSCs; AlphaV^+^CD200^+^CD45^-^6C3^-^CD105^-^Ter-119^-^Tie2^-^Thy^-^), expression of the following combination of transcripts was investigated in the *Grem1* AC population. Expression of *Itga5* (encoding AlphaV) and *Cd200* (encoding CD200) and lack of expression of *Ptprc* (encoding CD45), *Enpep* (encoding 6C3), and *Eng* (encoding CD105). Transcripts for *Ly76* (encoding Ter119), *Tek* (encoding Tie2) and *Thy1* (encoding Thy1.1/Thy1.2), were not detected in the AC populations and so were not included in the mSSC transcriptomic selection criteria. All violin plots were generated with VlnPlot() function in Seurat package and Chi-Square tests were performed with chisq.test() function in the base package.

**Statistical Analysis**

All analyses were performed using Prism 8/9 (GraphPad software Inc.), individual test details are provided in Figure legends. Line in dot plots indicate mean.

**KEY RESOURCES TABLE**

| REAGENT or RESOURCE | SOURCE | IDENTIFIER |
| --- | --- | --- |
| Antibodies | | |
| PCNA | Abcam | #ab18197 |
| COLX | Abcam | #ab28484 |
| COL2 | Abcam | #ab34712 |
| SOX9 | Millipore | #AB5535 |
| Anti-RFP | Rockland | #600-401-379 |
| Anti-Rabbit Biotin | VectorLab | #BA-1000 |
| Streptavidin-HRP | VectorLab | #SA-5004 |
| Anti-CD45 | Biolegend | #103131 |
| Ter119 | Biolegend | #116227 |
| Anti-CD31 | BD Pharmingen | #562861 |
| Anti-CD200 | Abcam | #ab33735 |
| APC/Cy7 Streptavidin | Biolegend | #405208 |
| DAPI | Sigma | #D9542 |
| DAB Chromogen | Dako | #K3468 |
| Pregnant Mare Serum Gonadotrophin | Intervet |  |
| human chorionic gonadotrophin | Intervet |  |
| FHM Media | Merck Milipore |  |
| KSOMMAA media | Merck MIlipore |  |
| Type VII collagenase | Sigma | #C0773 |
| Antigen unmasking solution | VectorLab | #H-3300 |
| Collagenase IV | Gibco | #17104019 |
| Dispase | Gibco | #17105041 |
| α-MEM | Sigma | #M5650 |
| Collagenase type I | Worthington | #CLS-1 |
| Collagenase type II | Worthington | #CLS-2 |
| Bacterial and virus strains | | |
| Biological samples |  |  |
| Chemicals, peptides, and recombinant proteins | | |
| Pregnant Mare Serum Gonadotrophin | Intervet |  |
| SpCas9 protein | PNA Bio |  |
| Tamoxifen | Sigma | #T5648 |
| Diphtheria toxin | Enzo | #BML-G135 |
| Osteosoft | Merck Milipore | #101728 |
| Cryofilm | Section-Lab | Type IIC(10) |
| Toluidine blue | Sigma | #198161 |
| Safranin O | Sigma | #8884 |
| Fast Green | Sigma | #F7252 |
| Alcian Blue | Sigma |  |
| Oil Red O | Sigma |  |
| Alizarin Red | Sigma |  |
| TRIzol | Thermofisher |  |
| Critical commercial assays | | |
| High Pure PCR template preparation kit | Roche |  |
| Click-iT EDU proliferation kit | Thermofisher | #C10086 |
| StemPro Osteogenic media |  |  |
| StemPro Chondrogenic media |  |  |
| StemPro Adipogenic media |  |  |
| SuperScript IV reverse transcriptase kit | Invitrogen |  |
| FastStart TaqMan Probe Master | Roche | #04673409001 |
| Deposited data | | |
| GEO |  | GSE193175 |
| Experimental models: Cell lines | | |
| Experimental models: Organisms/strains | | |
| *LepR-cre* | Jackson Laboratory | #008320 |
| *Acan-creER^T2^* | Jackson Laboratory | #019148 |
| *Grem1-creER^T^* | Jackson Laboratory | #027039 |
| *R26-LSL-TdTomato* | Jackson Laboratory | #007914 |
| *R26-LSL-ZsGreen* | Jackson Laboratory | #007906 |
| *R26-LSL-iDTR* | Jackson Laboratory | #007900 |
| Oligonucleotides | | |
| GAPDH | IDT probes | Mm.PT.39a.1 |
| Grem1 | IDT probes | Mm.PT.53a31803129 |
| Recombinant DNA | | |
| Software and algorithms | | |
| R environment |  | Version 4.0.2 |
| Seurat Package |  | Version 3.2.2 |
| GraphPad software Inc |  | Prism 8/9 |
| Flo Jo |  | v10.6.2 |
| ImageJ | NIH | v2.1.0/1.53d |
| Other | | |
| Clonal Cylinders | Sigma | #Z370789 |
